# Protein design of two-component tubular assemblies like cytoskeletons

**DOI:** 10.1101/2024.04.17.589732

**Authors:** Masahiro Noji, Yukihiko Sugita, Yosuke Yamazaki, Makito Miyazaki, Yuta Suzuki

**Affiliations:** Research Fellow of Japan Society for the Promotion of Science, Japan; Graduate School of Human and Environmental Studies, Kyoto University, Kyoto, Japan; Institute for Integrated Cell-Material Sciences, Kyoto University, Kyoto, Japan; Institute for Life and Medical Sciences, Kyoto University, Kyoto, Japan; Graduate School of Biostudies, Kyoto University, Kyoto, Japan; Hakubi Center for Advanced Research, Kyoto University, Kyoto, Japan; Graduate School of Science, Kyoto University, Kyoto, Japan; RIKEN Center for Biosystems Dynamics Research, Yokohama, Japan; PRESTO, JST, Saitama, Japan

## Abstract

Recent advances in protein design have ushered in an era of constructing intricate higher-order structures^1^. Despite this progress, orchestrating the assembly of diverse protein units into cohesive artificial structures akin to biological assembly systems, especially in tubular forms, remains an elusive goal. To address this, we introduce the Nature-Inspired Protein Assembly Design (NIPAD), a novel methodology that utilises two distinct protein units to create unique tubular structures under carefully designed conditions. These structures demonstrate dynamic flexibility similar to that of actin filaments, with cryo-electron microscopy revealing diverse morphologies, like microtubules. By mimicking actin filaments, helical conformations were incorporated into tubular assemblies, thereby enriching their structural diversity. Remarkably, these assemblies can be reversibly disassembled and reassembled in response to environmental stimuli, such as changes in salt concentration and temperature, mirroring the dynamic behaviour of natural systems. NIPAD combines rational protein design with biophysical insights, leading to the creation of biomimetic, adaptable, and reversible higher-order assemblies. This approach deepens our understanding of protein assembly design and complex biological structures. Concurrently, it broadens the horizons of synthetic biology and material science, holding significant implications for unravelling life’s fundamental processes and pioneering new applications.

## Main Text

Life phenomena rely on the dynamic and reversible assembly and disassembly of various higher-order protein assemblies. Actin filaments^2,3^ and microtubules^4,5^ in the cytoskeleton and the capsid proteins of viruses^6,7^ are examples of such naturally occurring structures. These are tightly regulated in function and complexity. Synthesizing higher-order structures of heterogeneous protein units poses a significant challenge, particularly in terms of replicating the diversity and flexibility inherent in natural assemblies. Although recent advances in computational design have enabled the creation of artificial higher-order protein structures from two protein components^8–10^, the design of heterogeneous higher-order protein assemblies with the flexibility and reversible assembly/disassembly characteristics of natural structures, especially tube structures reminiscent of the cytoskeleton, remains a formidable challenge.

Herein, we introduced Nature-Inspired Protein Assembly Design (NIPAD), a novel methodology that draws inspiration from the principles underlying natural protein complexes. By integrating rational protein design with biophysical insights to optimise assembly conditions, NIPAD recapitulates flexibility and reversible assembly principles. We employed NIPAD to create a novel assembly of two distinct protein units, successfully forming unique two-component tube structures. This development represents a significant step towards replicating the properties of complex natural structures at the molecular level.

### The concept of NIPAD

In developing the protein components for NIPAD, we employed rational design principles with hints from natural biological systems, integrating naturally occurring ’heterolinkers’ with ’scaffold proteins’ to streamline design. For the heterolinker, we chose the heterodimeric peptide pair “MBD3L2 (M3L2)/p66α” (**Fig. 1a**). Our choice of M3L2/p66α was influenced by its role in the MBD2-NuRD complex, where the ’MBD2/p66α’ anti-parallel coiled-coil domain is essential for complex assembly^11^. Given the moderate denaturation midpoint temperature (*T*_m_) of M3L2/p66α (*T*_m_ = 35 °C) compared to MBD2/p66α (*T*_m_ = 65 °C)^12^, we anticipated that M3L2/p66α would provide a balance between stability and reversible assembly control through temperature modulation. We then sought to identify a scaffold protein that could connect the heterolinker in the simplest manner possible. Previous study has shown that the positions of connecting sites at the corners of such scaffold proteins can facilitate desired assembly formation^13^. Therefore, we chose the “*Pseudomonas fluorescens* PuuE allantoinase (PuuE)”, a homotetramer with *C*_4_ symmetry where each C-terminus is located at each vertex of the quaternary structure (**Fig. 1b**)^14^. This arrangement enabled straightforward genetic fusion of heterolinkers to the scaffold’s C-termini, leveraging specificity and reversibility of heterolinker interactions to drive assembly formation. This approach not only simplifies the assembly process but also enhances expression and purification efficiency for each protein unit, preventing spontaneous assembly and ensuring the controlled formation of higher-order structures.

**Figure 1.**
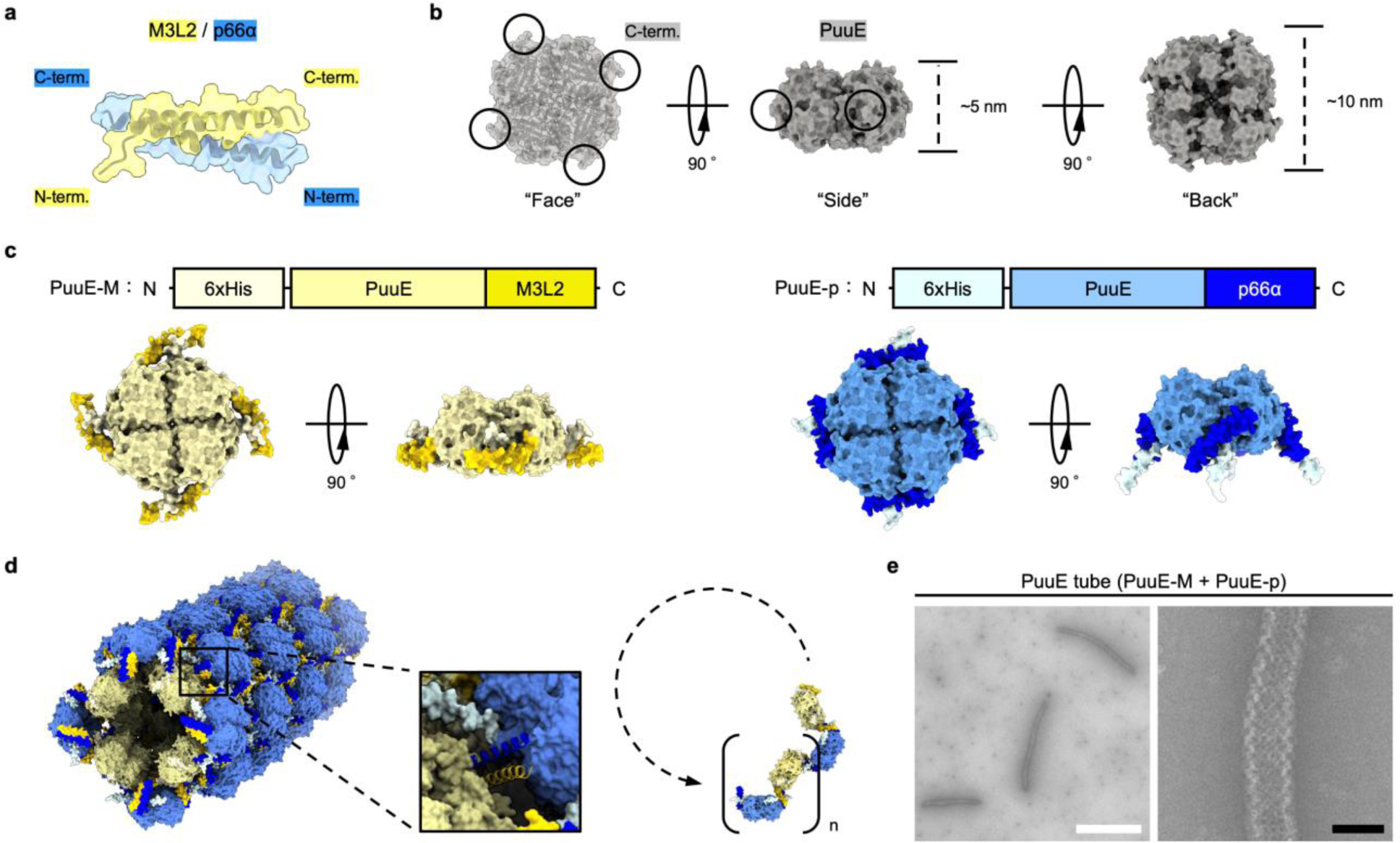
Construction of a PuuE tube via NIPAD. **a**, AF2 prediction of the heterodimeric peptide pair, M3L2 (yellow) and p66α (blue). **b,** Crystal structure of PuuE (PDB: 3CL6). C-terminus positions are circled. Detailed structure, face, side, and back are shown for clarity. **c,** Schematic diagram of the protein sequence (top) and the AF2 predicted structures of PuuE-M and PuuE-p (bottom). PuuE-M and PuuE-p are coloured yellow and blue to match the respective peptides and overall structure to clear the tube structure (**d**). The peptide parts, M3L2 and p66α, are highlighted in darker colours. **d,** Left, predicted model of the tubular assembly consisting of PuuE-M and PuuE-p. Right, brief schematic diagram of how many proteins (n) form a system of tube structures. **e,** nsTEM images of tubular assemblies constructed from PuuE-M and PuuE-p. 12.5 μM PuuE-M and 12.5 μM PuuE-p in NaCl (+) buffer was incubated at 40 °C for 24 h and imaged via nsTEM. Scale bars, 1 μm (white), 100 nm (black).

We constructed protein units ’PuuE-M’ and ’PuuE-p’ through genetic engineering, fusing M3L2 and p66α to the C-terminus of PuuE, respectively (**Fig. 1c**). AlphaFold2 (AF2)^15,16^ modelling suggested a configuration with a relatively flexible orientation of M3L2 in PuuE-M, whereas a highly constrained orientation of p66α in PuuE-p (**Extended Data Fig. 1**). Due to the constrained orientation of PuuE-p, an angular interface was formed between PuuE-M and PuuE-p, and we predicted that assembly of these two units would form a tubular structure (**Fig. 1d**). In addition, depending on the number of PuuE-M and PuuE-p units, different tubular structures were expected. Protein expression in *Escherichia coli* (*E. coli*) provided both constructs in a soluble form, facilitating their purification. In isolation, neither protein unit exhibits self-assembly (**Extended Data Fig. 2a, b**). However, when combined under optimised conditions (discussed in the following section), we successfully observed the expected chessboard-patterned tube (PuuE tube) using negative-stain transmission electron microscopy (nsTEM) (**Fig. 1e**, **Extended Data Fig. 2c**). Although previous studies have assembled cages^8,9^, sheets^10^, and three-dimensional (3D) crystals^17^ using two-component protein systems, this study is unique in that tube structures were successfully created.

### The condition design of tubular assemblies

Based on established principles observed in biological systems such as actin filaments^18,19^, microtubules^20,21^, and amyloid fibrils^22,23^, protein concentration, temperature, time, and salinity have a significant influence on assembly formation. Thus, we carefully tailored assembly conditions to exploit the complex interactions between these factors. This approach allowed us to optimise experimental conditions for constructing the desired tubular structures.

First, we focused on the dependency of PuuE tube assembly on protein concentration (**Extended Data Fig. 3**). Mixing PuuE-M and PuuE-p at a concentration of 250 nM each (considering tetramer equivalence) led to the formation of tubular structures after an incubation period of 24 h at 40 °C, consistent with the dissociation constant (*K*_d_) for M3L2/p66α dimer formation, which is approximately 268 nM^12^. Increasing protein concentration to 2.5 µM markedly enhanced the quantity and length of formed tubular structures. Elevating the concentration to 12.5 µM for each component significantly increased tube formation efficiency, underscoring the concentration-dependent nature of PuuE-M- and PuuE-p-facilitated tubular assembly.

Next, PuuE tube formation kinetics were investigated. The incubation of mixtures containing 12.5 µM of each protein at 40 °C resulted in the formation of nascent tube structures within 30 min, evolving into distinguishable tubes spanning several hundred nanometres to 1 μm in length within 1-2 h. Over time, these tubes elongated, reaching several micrometres in length after 24 h and extending up to approximately 5 μm after 48 h (**Fig. 2a, b**, **Extended Data Fig. 4a**). Once formed, the tubes remained structurally stable for at least one month at room temperature (**Extended Data Fig. 4b**).

**Figure 2.**
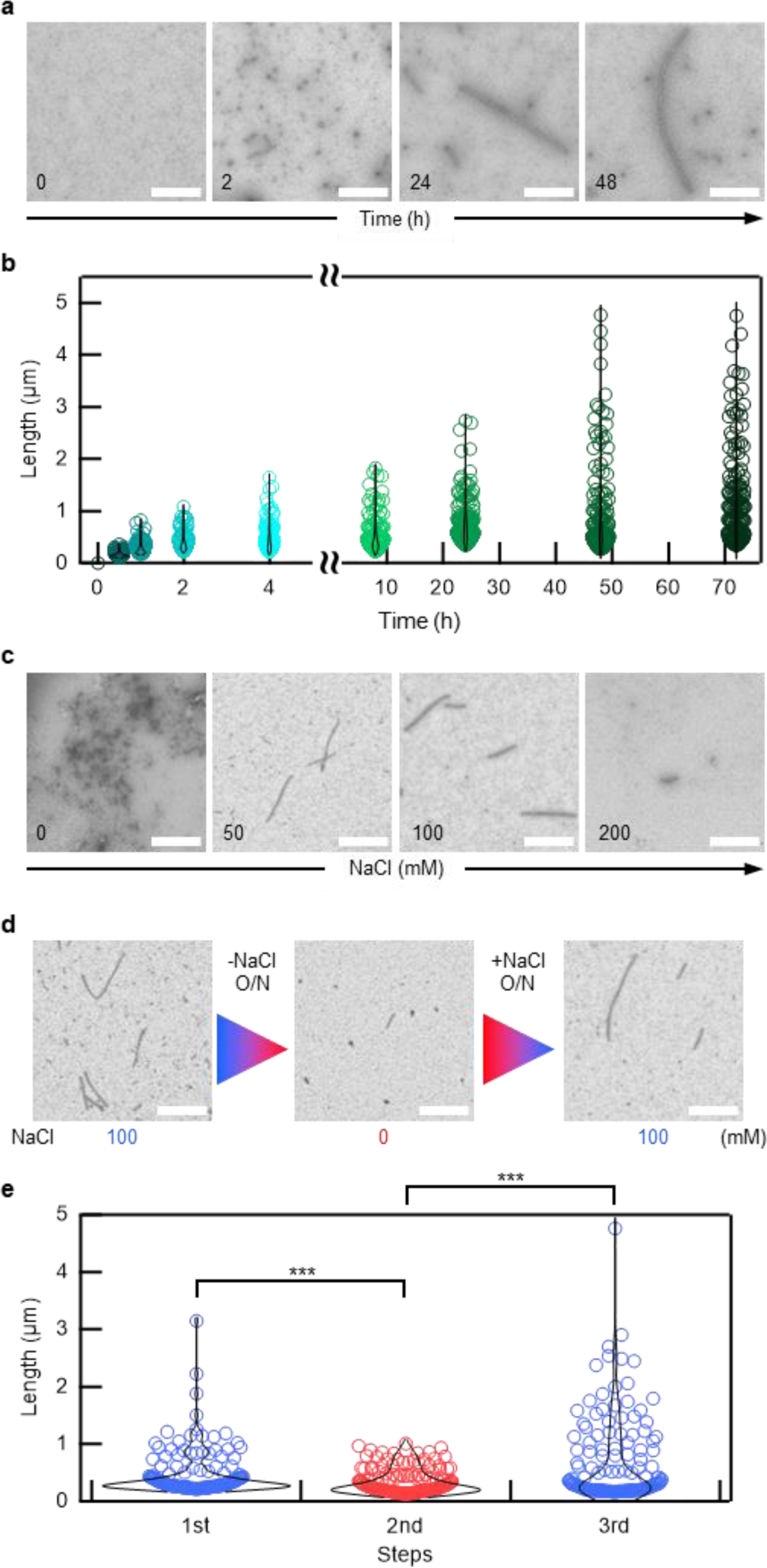
Condition optimisation for PuuE tube assembly. **a, b**, The kinetics of tubular assembly. nsTEM images of tubular assembly (**a**) and length analysis (**b**). **c,** nsTEM images of tubular assemblies with varying NaCl concentration. **d,** nsTEM images showing the reversibility of tube structures with changing NaCl concentration. **e,** Tube length analysis of nsTEM images. For tube length analysis, tubes were picked up and calculated from 5k images at each step. 150 tubes from the longest tube length were used at each data point. *** p<0.001 (Welch’s t-test). Scale bar, 1 µm.

We then explored the influence of temperature on PuuE tube formation (**Extended Data Fig. 5a**). While the melting temperature of the M3L2/p66α dimer is around 35 °C, tube assembly was hardly observed at sufficiently lower temperatures of 20-25 °C, even after 24 h of incubation. In contrast, temperatures near *T*_m_, specifically between 30 and 40 °C, markedly promoted tube formation. This observation suggests that temperatures below *T*_m_ excessively enhance the binding force between M3L2 and p66α, causing kinetic entrapment of assemblies. However, temperatures close to *T*_m_ modulate this binding force, allowing the dynamic rearrangement of M3L2/p66α interactions under thermal fluctuations, thus facilitating assembly of thermodynamically stable, ordered structures. This principle is consistent with general crystallisation theories^24,25^ and reports on the formation of ordered structures in natural protein assemblies^22,23,26^. It should be noted that temperatures above 45 °C led to thermal denaturation and aggregation of PuuE-M (*T*_m_ = 46.2 °C) and PuuE-p (*T*_m_ = 48.1 °C), significantly diminishing tube formation capabilities (**Extended Data Fig. 5b, c**). This implies that the original concept of tube formation with reversible temperature control was not realised.

### Reversibility of tubular assemblies

Finally, we examined the effects of salt concentration on PuuE tube assembly. We prepared mixtures in varying NaCl concentrations ranging from 0 to 400 mM and incubated them at 40 °C for 24 h. Tube formation was clearly observed within the NaCl concentration window of 50-200 mM, with no tube formation detected outside this range (**Fig. 2c**, **Extended Data Fig. 6a**). Since both PuuE-M (pI = 6.44) and PuuE-p (pI = 6.02) were similarly charged under pH 8.0, tube formation at low salt concentrations was likely inhibited by electrostatic repulsion. Conversely, moderate electrostatic shielding facilitated by 50-200 mM NaCl likely provided conducive conditions for tube assembly, whereas higher NaCl concentrations may have induced excessive shielding or aggregation due to salting out, inhibiting tube formation. This observation aligns with known phenomena in protein crystallisation, where electrostatic shielding above a certain threshold can prevent crystal growth^17,27–29^, although crystals formed by a combination of electrostatic and hydrophobic interactions can remain stable up to approximately 200 mM NaCl^30^. The association of M3L2/p66α involves both electrostatic and hydrophobic interactions^12^, consistent with the latter scenario.

The salt-dependent PuuE tube formation and the dynamic nature of PuuE-M/PuuE-p interactions near their *T*_m_ (35 °C) led us to hypothesise that tubes could undergo reversible disassembly and reassembly in response to changes in NaCl concentration. Confirming our hypothesis, tubes initially formed in 100 mM NaCl solution were significantly shortened when subjected to solvent exchange with a 0 mM NaCl buffer (NaCl (-) buffer) and subsequent incubation at 40 °C for 24 h (**Fig. 2d, e**, **Extended Data Fig. 6b**). Subsequent solvent exchange with 100 mM NaCl buffer (NaCl (+) buffer) resulted in notable tube reassembly. This salt-concentration-driven reversibility, although divergent from the initial temperature-controlled reversibility hypothesis, marks a significant advance in artificial protein assembly design, allowing for the biomimetic replication of dynamic structural changes under relatively mild conditions, akin to the behaviour of actin filaments in cellular structures^18,19^. Unlike irreversible aggregation observed in amyloid structures, our assemblies exhibit a reversible and dynamic assembly process akin to the behaviour of the cytoskeleton, successfully demonstrating the potential for biomimetic replication of natural cellular dynamics under controlled conditions.

### Diversity and flexibility of tubes

Based on these findings, we determined the ideal conditions for PuuE tube formation in 100 mM NaCl at 40 °C for 24 h. To further characterise the structural features of tubes formed under these conditions, cryo-electron microscopy (cryo-EM) was employed (**Fig. 3a**, **Extended Data Fig. 7a**). Analysis of 2D class-averaged images revealed a spectrum of tube diameters and symmetries similar to the diversity observed in microtubules^31–34^ (**Fig. 3b**). From these images, we successfully reconstructed the 3D structures with *C*_4_, *C*_5_, and *C*_6_ symmetries within tube structures (**Fig. 3c**). Insights from the PuuE crystal structure^14^, notably its unique central indentation on the back surface (**Fig 1b**), allowed us to deduce that PuuE units are alternately oriented face-to-back across all 3D models. In addition, cryo-EM analysis suggested that the flexibility of connections allowed contraction of the entire tube structure (**Extended Data Fig. 7b**). Due to the flexibility, the resolution was insufficient to differentiate between PuuE-M and PuuE-p units. Furthermore, tubes with larger diameters, presumably having *C*_7_ to *C*_10_ symmetries, were identified at low resolution, likely due to the flexibility of connection sites influencing tube structure. In fact, nsTEM and cryo-EM images frequently showed tubes appearing bent or compressed (**Extended Data Fig. 2–7**). In contrast to prior strategies yielding relatively uniform higher-order protein assemblies by integrating binding sites at specific points^13,35^ or surfaces^8–10^ on scaffold proteins, NIPAD integrates a flexible linker with the scaffold protein, resulting in varied structures and arrangements among higher-order assemblies. This variation in tube diameter, akin to that observed in microtubules^31–34^, is presumably a hallmark of NIPAD.

**Figure 3.**
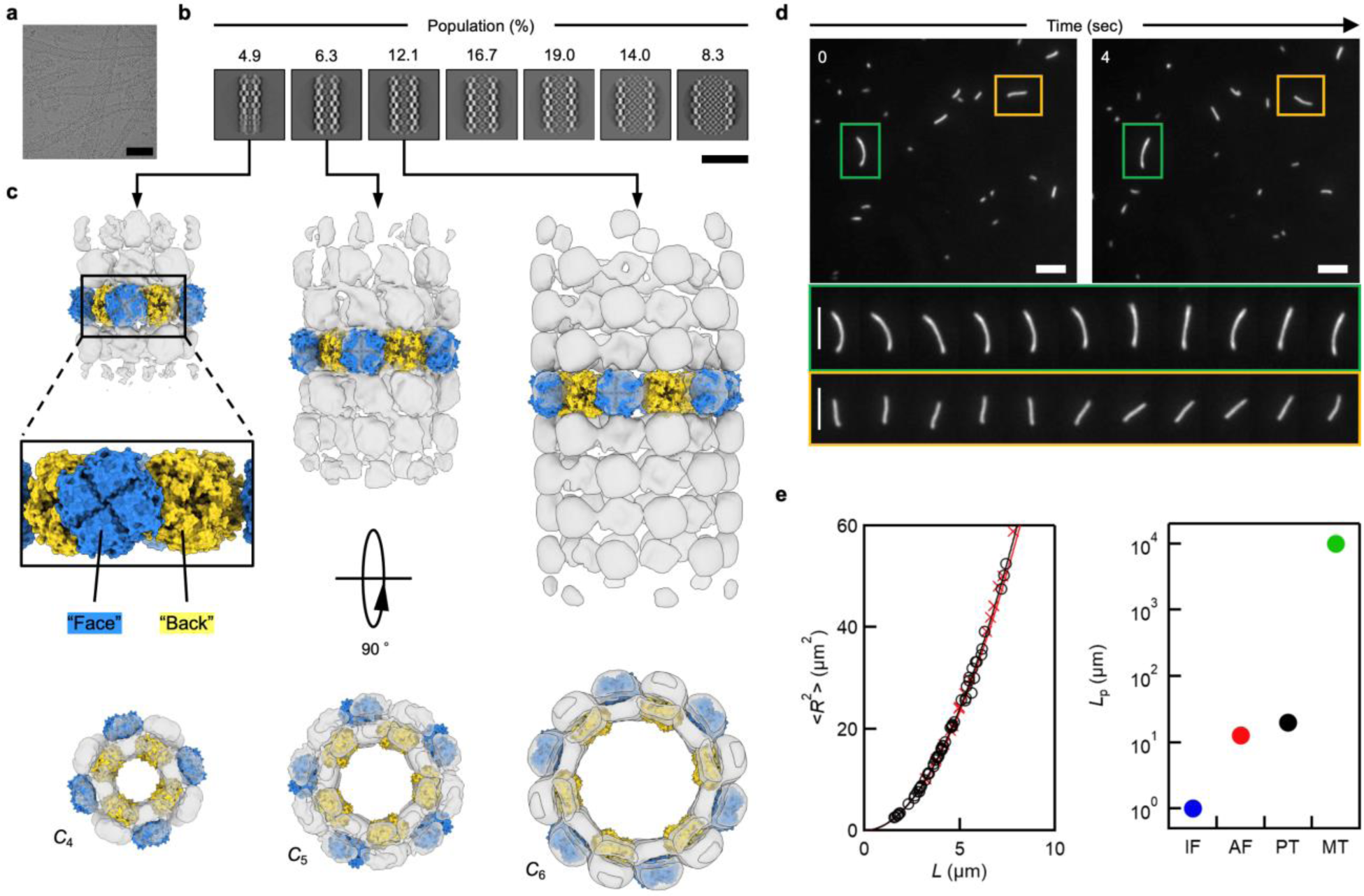
Structural characterisation of PuuE tube. **a**, Representative cryo-EM image of tube structures. **b,** 2D class-averaged images of tube structures. The population of each structure was determined from the total pickings of 206,658 tube segments. Scale bar, 500 Å. **c,** 3D reconstructed models of tube structures with *C*_4_, *C*_5_, and *C*_6_ symmetries. Due to flexibility of tube structure, PuuE-M and PuuE-p were indistinguishable, but colour-corded for clarification. **d,** Time-lapse images of random bending of the tube structures observed via TIRFM. Top: snapshots at starting point (0 sec) and after 4 sec (top). Bottom: enlarged images of tubes in green or orange rectangles in the top images, showing the dynamic flexibility of tube structures between 0 to 4 sec (0.4 sec per image). Scale bar, 5 μm. **e,** Left, a relationship between contour length (*L*) and mean square of end-to-end distance (<*R*^2^>) of the tube structures for estimation of the persistence length (*L*_p_). The continued lines represent fitting curves (black for PuuE tube, red for actin filament) to experimental data (black open circle for PuuE tube, red cross mark for actin filament). Right, comparison of persistence length with cytoskeletal elements. PuuE tube (PT, black) and actin filaments (AF, red) were determined in this study (A wider range of plots is shown in an **Extended Data Fig. 9**). Intermediate filaments (IF, blue) and microtubules (MT, green) are taken from ref. 41 and 38, respectively.

To further explore PuuE tube structure flexibility, we labelled tubes with Alexa Fluor 488 succinimidyl ester and observed them in real-time via total internal reflection fluorescence microscopy (TIRFM). Tube structures were constrained in the evanescent field by the depletion effect of methylcellulose contained in the observation buffer and underwent thermally driven two-dimensional random bending (**Fig. 3d**, **Extended data Fig. 8a**). Analysis of the fluctuation in shape yielded the persistence length (*L*_p_) of 19.7 µm (**Fig. 3e**, **Extended data Fig. 8b**). *L*_p_ is the mean length over which a semiflexible polymer remains straight, characterising polymer stiffness^36^. The *L*_p_ value of the tube structures is nearly equal to that of actin filaments measured in this study, 12.5 µm (**Fig. 3e**), and previously reported values of 9-20 µm^37^. Microtubules have much longer persistence lengths, ranging from 0.1 to 10 mm^38–40^. In contrast, intermediate filaments, another cytoskeletal fibre structure, typically have shorter persistence lengths; < 1 µm^41^. Therefore, the tube structure is as flexible as actin filaments, more flexible than microtubules, and stiffer than intermediate filaments.

### Emulation of actin filaments

Finally, we sought to modify the morphology of PuuE tube assemblies. Specifically, we hypothesised that grafting the D-loop of actin onto PuuE-M would produce tubes with a helical conformation reminiscent of actin filaments. The D-loop plays an important role in helical actin filament formation via hydrophobic pockets^42–45^. The hydrophobic nature of a prominent indentation on the “back” side of PuuE (**Extended Data Fig. 9a**) guided our hypothesis. The loop structure on the back side of PuuE-M was chosen as the grafting site for the D-loop, and the ’PuuE(D-loop)-M’ fusion construct was constructed (**Fig. 4a**). When PuuE(D-loop)-M was expressed in *E. coli*, it was found in the soluble fraction and was purified as PuuE-M. Although PuuE(D-loop)-M alone did not assemble, its combination with PuuE-p not only replicated the PuuE tube but also introduced novel helical patterns, with two or three tubes intertwined (PuuE D-loop tube), as verified via nsTEM (**Fig. 4b**, **Extended Data Fig. 9b**). While the D-loop has been suggested to play a crucial role in the helical formation of actin filaments^42–45^, its complete mechanism remains not fully elucidated. Our study, by successfully grafting the D-loop to replicate actin-like helical structures, offers a novel perspective on the significance of the D-loop. This approach not only confirms the critical role of the D-loop in helical conformations but also opens new avenues for understanding the intricate design principles of actin filaments.

**Figure 4.**
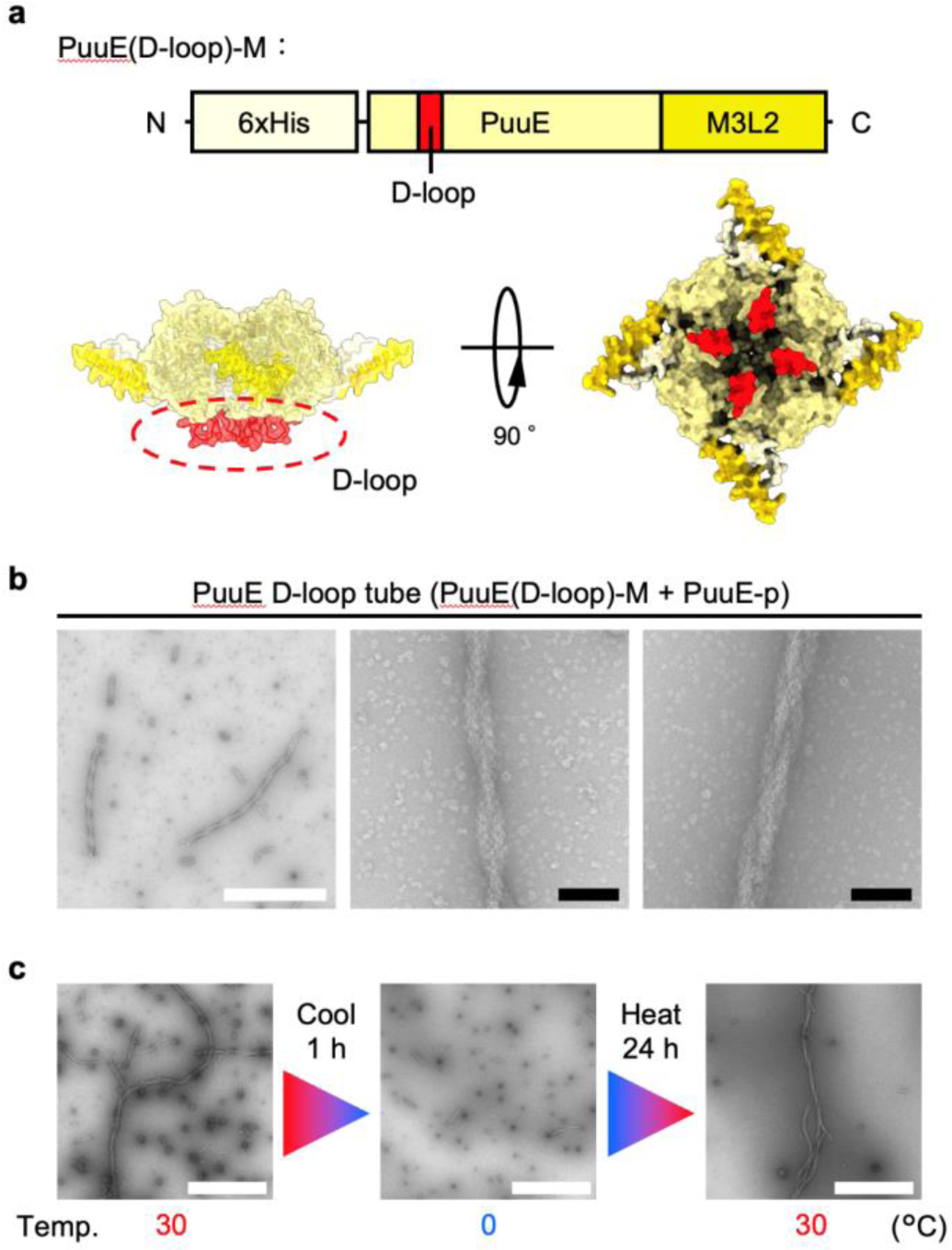
Emulation of actin filament by D-loop grafting. **a**, Schematic representations of PuuE(D-loop)-M. The position of D-loop graft (red) is indicated by protein sequence (top) and the AF2 predicted structure (bottom). **b,** nsTEM images of tubes with a helical conformation composed of PuuE(D-loop)-M and PuuE-p. The helical pattern of two (centre) or three (right) intertwined tubes was shown in the high magnification image. **c,** nsTEM images showing the reversibility of tube structure with helical conformations by temperature change. Scale bars, 1 μm (white), 100 nm (black).

Since PuuE(D-loop)-M has a lower thermal stability (*T*_m_ = 35.6 °C) than PuuE-M (*T*_m_ = 46.2 °C, **Extended Data Fig. 5b, 9c**), we had to incubate at a lower temperature (30 °C). The emergence of helical formations, absent in the PuuE-M and PuuE-p mixtures, clearly stems from D-loop integration. This enriches our understanding of the D-loop contribution to the helical architecture of actin filaments and demonstrates the effectiveness of bottom-up approaches in elucidating protein structural and functional dynamics. As mentioned above, the helical conformation of tube structures is thought to arise from hydrophobic interactions, which are inherently sensitive to temperature and weaken at lower temperatures^46–48^. This led us to posit that alterations in temperature can serve as reversible switches for disassembly and reassembly. Intriguingly, exposing the samples to 0 °C for 1 h not only suggested a dissociation of the helical arrangements but hinted at a possible breakdown of the tubular structures (**Fig. 4c**, **Extended data Fig. 10**). Remarkably, when these disassembled samples were reintroduced to 30 °C for 24 h, the elongated tubular formations with helical conformations were restored. By grafting D-loop, the tube structure could not only form helical conformations but also acquired a new temperature-dependent reversibility. This thermal responsiveness parallels the behaviour of microtubules^20,21,49^, underscoring the ability of the NIPAD approach to mimic the dynamic properties of biomolecular assemblies in artificial protein design to create complex higher-order protein structures. This dual responsiveness (salt and temperature dependence) enhances the biomimetic potential of our design, which is a promising avenue for advanced applications in synthetic biology and materials science.

## Discussion

We introduce NIPAD, a pioneering approach that intricately weaves together protein unit design and assembly, drawing inspiration from the complexity and adaptability of natural protein assemblies. By employing NIPAD, we created a unique higher-order tubular assembly composed of two protein units, exhibiting the reversible, flexible, and diverse characteristics of natural structures. A noteworthy highlight of our study was the successful induction of helical conformations within these tube assemblies, akin to those observed in actin filaments, achieved through strategic integration of the D-loop into assembly design.

This advance in protein assembly highlights the complexity of emulating the dynamic behaviour observed in biological systems. The design and assembly of protein structures in vitro, although closely controlled, cannot fully replicate the complex cellular environment. In vivo, myriad factors such as macromolecular crowding, post-translational modifications and interactions with other cellular components can significantly influence protein behaviour^50^. Our designed protein assemblies exhibit remarkable biomimicry in terms of flexibility, reversibility, and structural diversity, but have yet to be demonstrated and validated in biological systems, where the true complexity of biological interactions is demonstrated. Furthermore, our approach, which focuses on the assembly of tubular structures inspired by cytoskeletal elements such as actin filaments and microtubules, does not address the full range of protein complex structures found within biological systems. Natural protein assemblies contain not only structural but also functional diversity, and there is much room for exploration beyond the scope of this study. Computational methods have an important role to play in improving the accuracy and breadth of protein assembly design^1^. By utilising computational predictions about protein interactions and assembly outcomes, our design would be refined into more complex and functional biomimetic structures, with applications ranging from novel biomaterials and nanodevices to therapeutic innovations.

Our research extends the boundaries of protein assembly design and provides new insights into its applications in synthetic biology and life sciences. This research encourages a comprehensive approach that bridges the divide between the biological and materials sciences, and suggests that the exploration of nature’s complex systems has the potential to transform science and technology. As we continue to explore this intersection of life and materials sciences, we anticipate that future investigations will provide fundamental insights into the natural world, heralding a new era of scientific discoveries and technological breakthroughs.

**Extended Data Figure 1.**
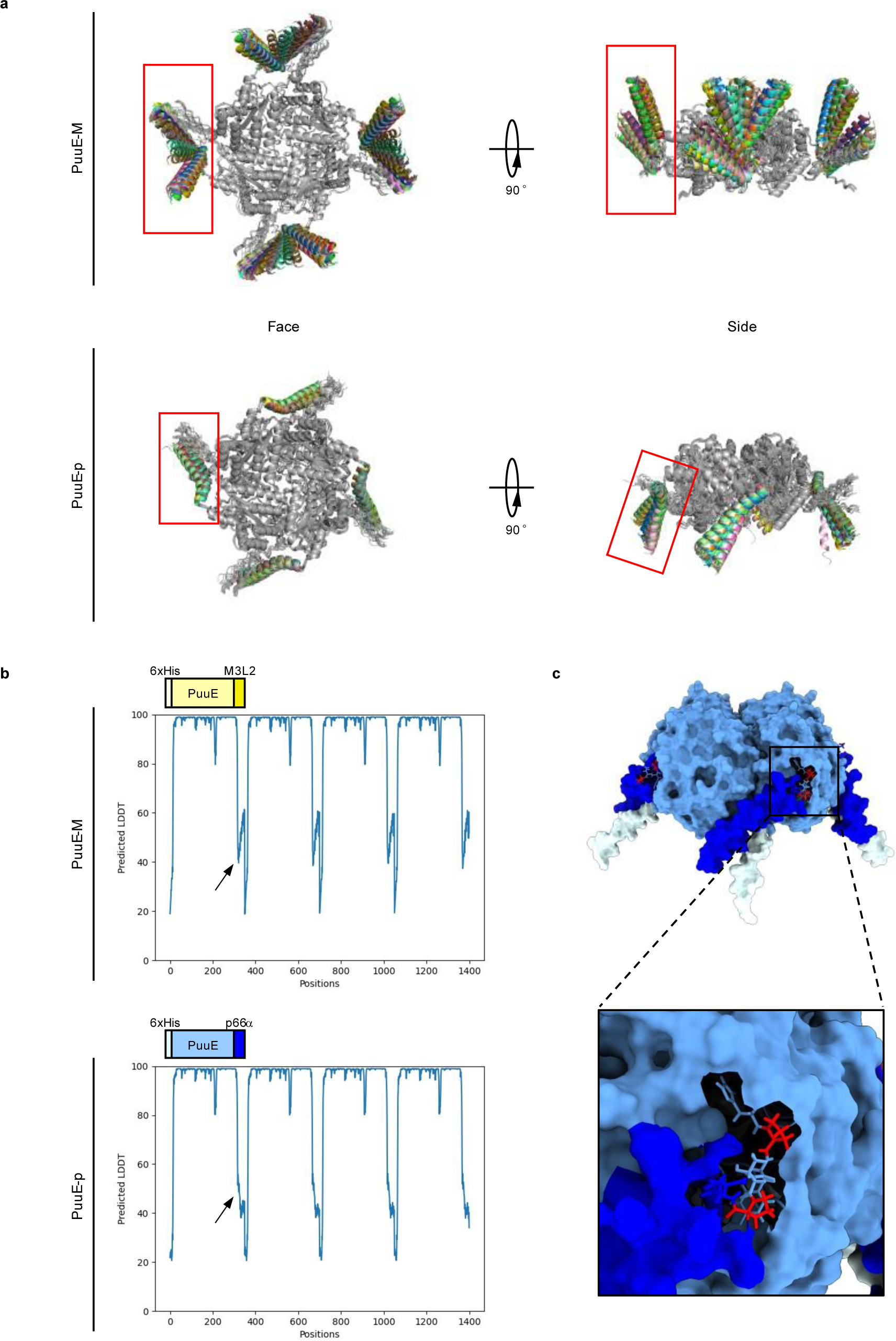
|AF2 prediction of PuuE-M and PuuE-p. **a**, Fifty prediction models overlapped for PuuE-M (top) and PuuE-p (bottom). Peptide parts, M3L2 and p66α, are circled in red box. **b,** Predicted local distance difference test plots for the most reliable prediction models for PuuE-M (top) and PuuE-p (bottom). Arrows indicate the N-terminal region of M3L2 and p66α. For these regions, PuuE-M has a lower predictive reliability than PuuE-p, suggesting that the structure may be more flexible. **c,** The most reliable prediction model for PuuE-p. The region from the C-terminus of PuuE to the N-terminus of p66α (i.e. ^313^HPYTPE^318^) is depicted by a stick model. The two Pro residues highlighted in red are thought to be responsible for the rigidity of the PuuE-p structure. Because of the rigidity of PuuE-p, the final product of the mixture was predicted to be a tube, as shown in Fig. 1d.

**Extended Data Figure 2.**
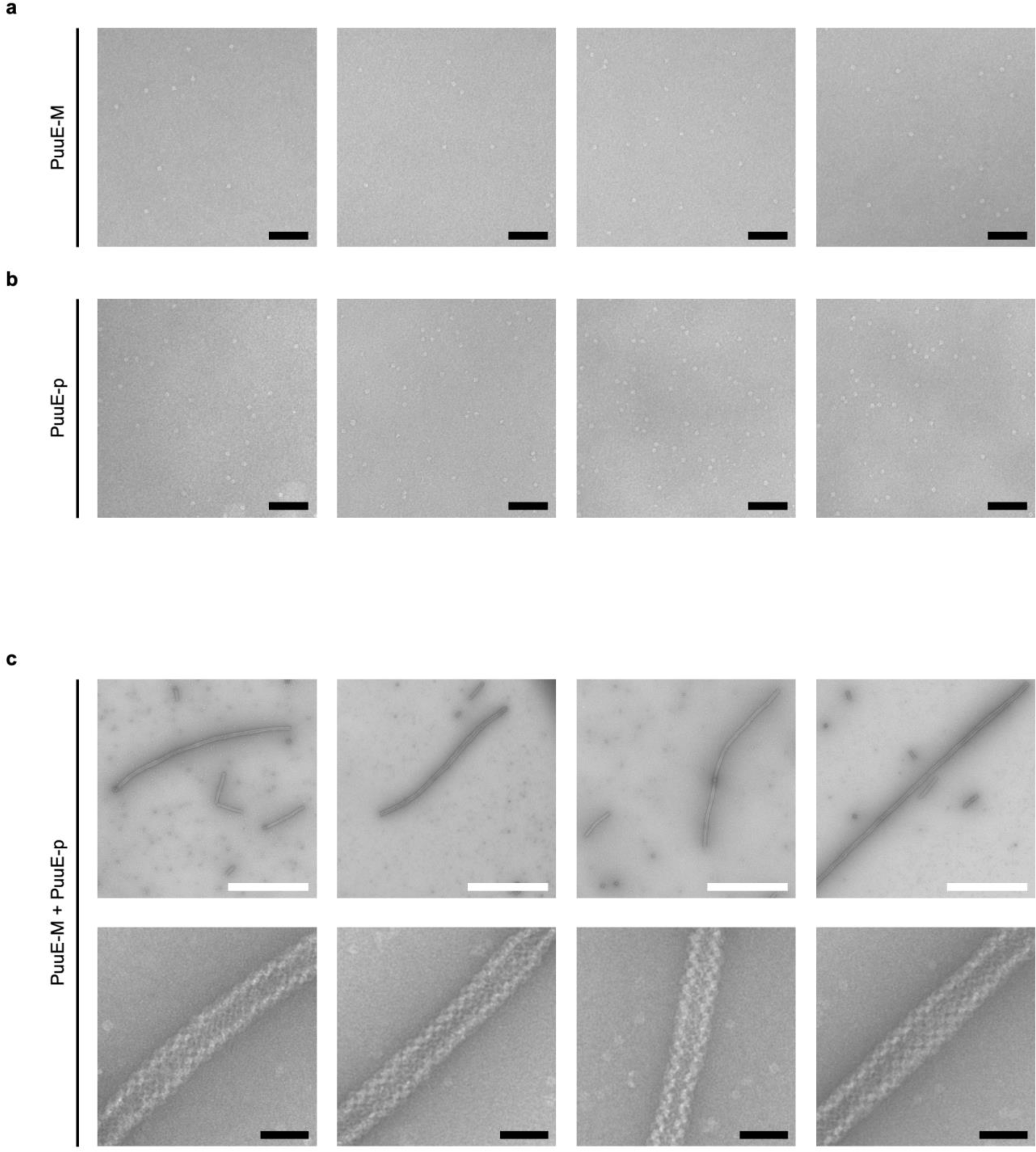
|nsTEM characterisation of PuuE-M, PuuE-p, and the mixture for PuuE-M and PuuE-p. **a**, 12.5 μM PuuE-M, **b,** 12.5 μM PuuE-p, or **c,** 12.5 μM PuuE-M and 12.5 μM PuuE-p in NaCl (+) buffer was incubated at 40 °C for 24 h (optimised conditions). The tube structure observed in the nsTEM images was flexible as it was curved and collapsed. Scale bars, 1 μm (white), 100 nm (black).

**Extended Data Figure 3.**
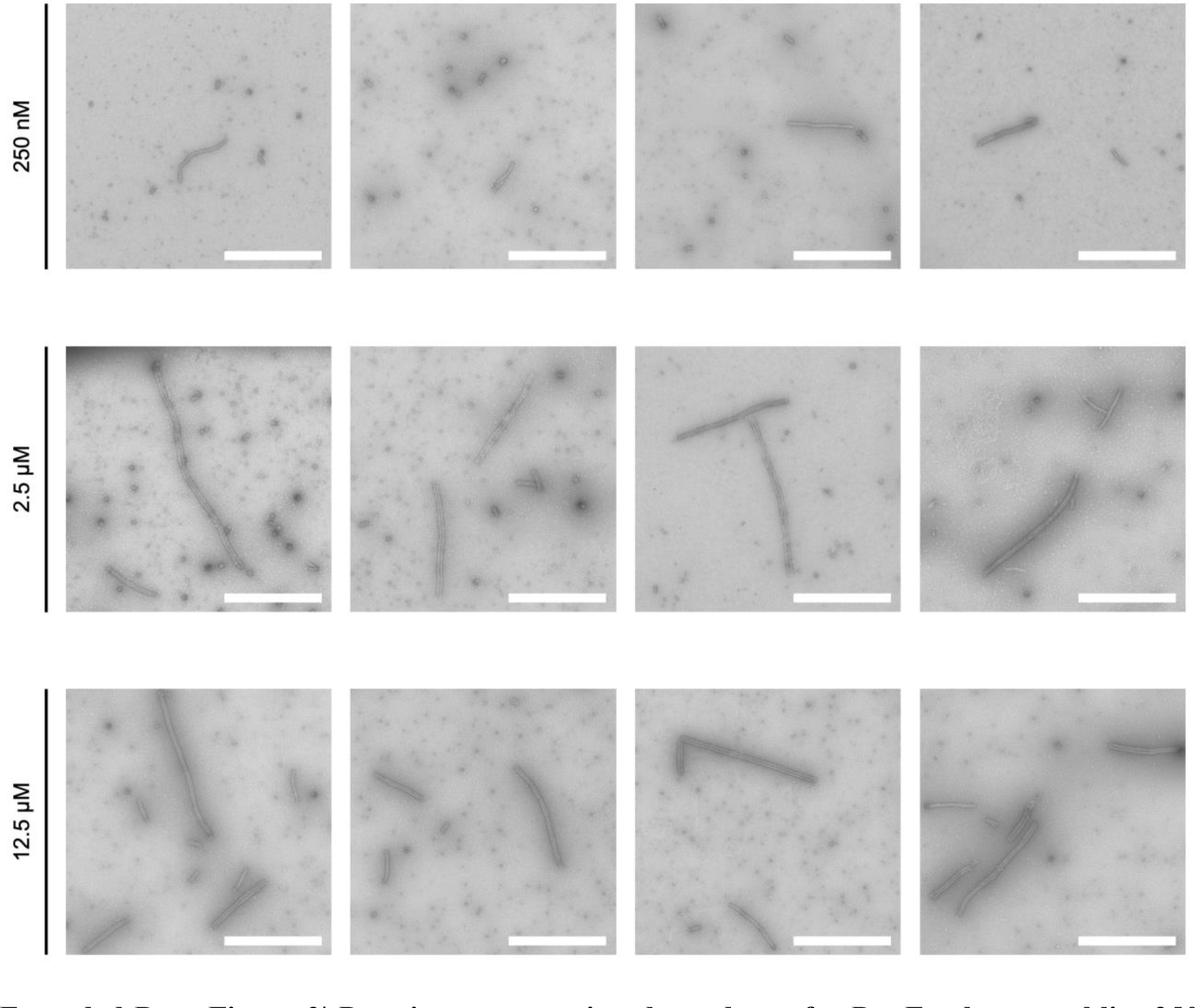
|Protein concentration dependence for PuuE tube assemblies. 250 nM (top), 2.5 μM (middle), and 12.5 μM (bottom) of PuuE-M and PuuE-p each in NaCl (+) buffer was incubated at 40 °C for 24 h and imaged by nsTEM. Scale bars, 1 μm.

**Extended Data Figure 4.**
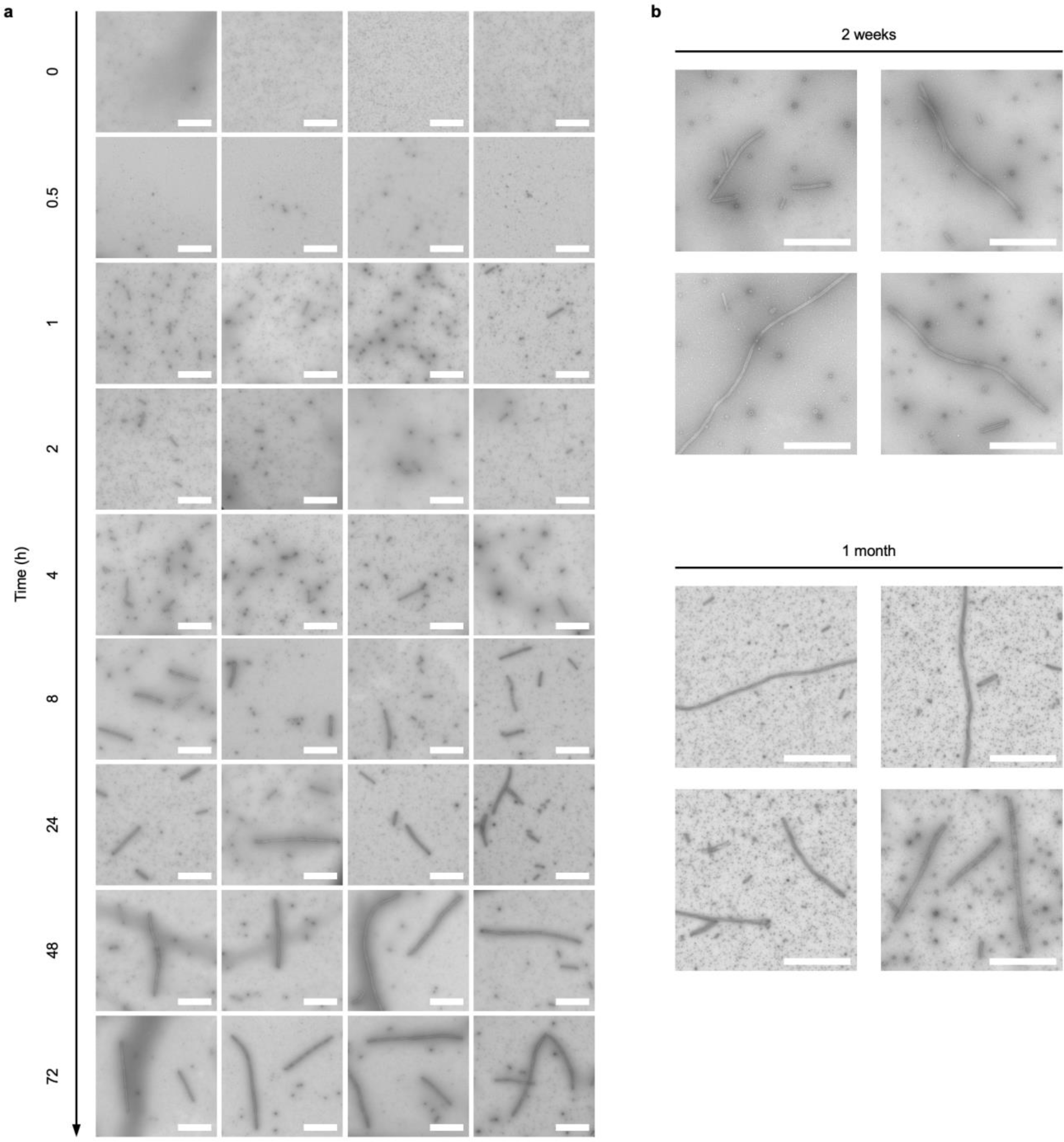
|Time dependence of PuuE tube assemblies and their stability over time. **a**, 12.5 μM of PuuE-M and PuuE-p each in NaCl (+) buffer was incubated at 40 °C for indicated time points and imaged via nsTEM. **b,** After 24 h of tube formation, the sample was kept at room temperature for the indicated time and imaged using nsTEM. Tube structures remained unchanged after 2 weeks and even after 1 month, suggesting stability. Scale bars, 1 μm.

**Extended Data Figure 5.**
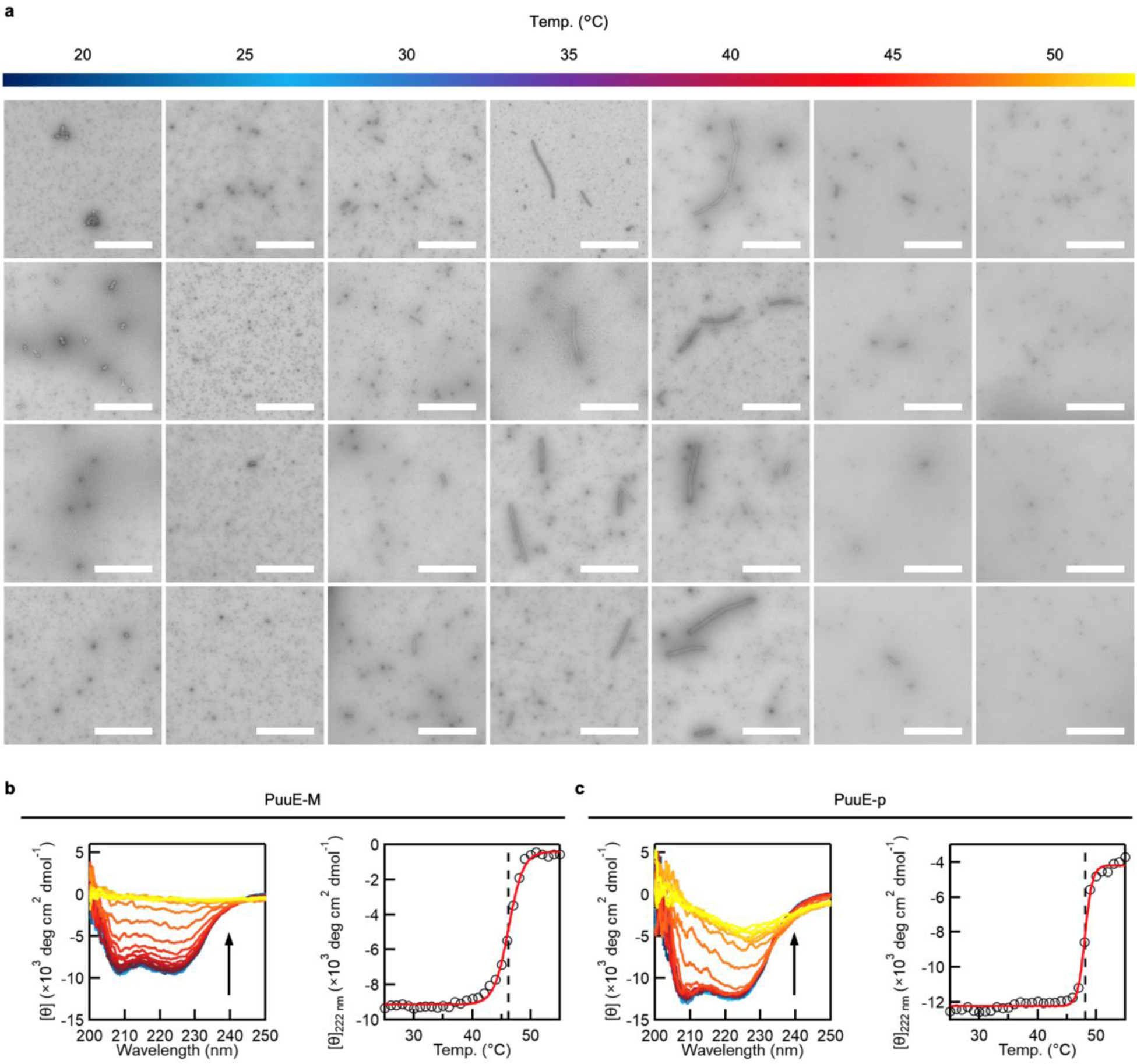
|Temperature dependence of PuuE tube assemblies and determination of *T*_m_ for PuuE-M and PuuE-p. **a**, 12.5 μM of PuuE-M and PuuE-p each in NaCl (+) buffer was incubated at indicated temperature for 24 h and imaged via nsTEM. Scale bars, 1 μm. **b, c,** *T*_m_ measurements using CD for PuuE-M (**b**) and PuuE-p (**c**). 2.5 μM of PuuE-M or PuuE-p in NaCl (+) buffer was incubated from 25 to 55 °C with temperature change of 1 °C/min. Left panel, overall CD spectra; right panel, thermal denaturation profiles.

**Extended Data Figure 6.**
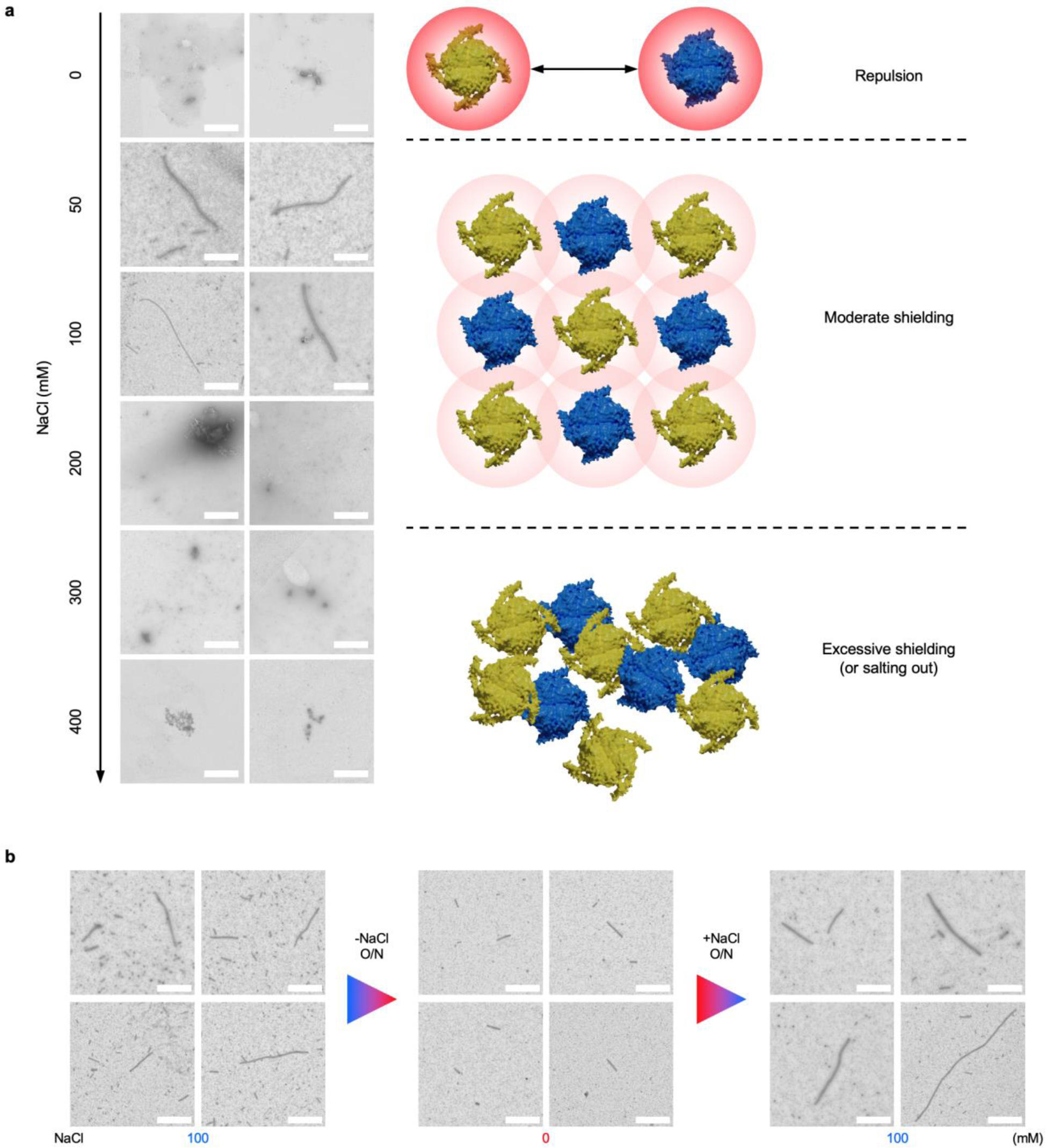
|Salt concentration dependence and reversibility of PuuE tube assemblies. **a**, Left, 12.5 μM of PuuE-M and PuuE-p each in NaCl (+) buffer was incubated at 40 °C for 24 h with indicated NaCl concentration and imaged via nsTEM. Right: diagram of salt concentration effects described in the main text. **b,** Additional images in Fig. 2d prove the reversibility of tubular assemblies. These images were used for statistical analysis of tube length, as shown in Fig. 2e. Scale bars, 1 μm (white).

**Extended Data Figure 7.**
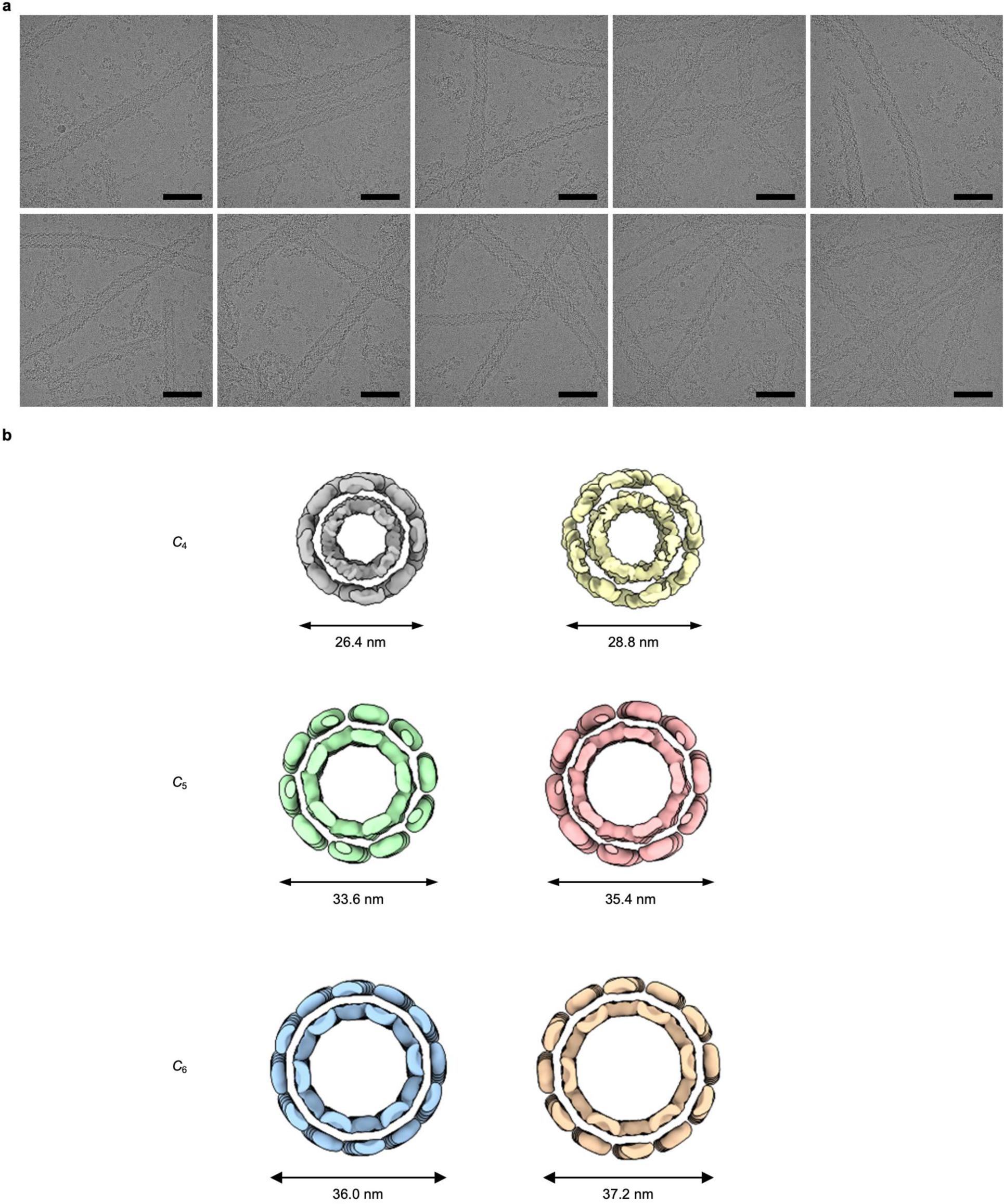
|Cryo-EM images and contraction of PuuE tube. **a**, Additional cryo-EM images of Fig. 3a. Different tube diameters and flexibility of the tube structures (curved and collapsed) were observed. Scale bars, 100 nm. **b,** Different diameters of PuuE tubes with *C*_4_, *C*_5_, *C*_6_ symmetry were observed from cryo-EM analysis, suggesting contraction of the PuuE tube.

**Extended Data Figure 8.**
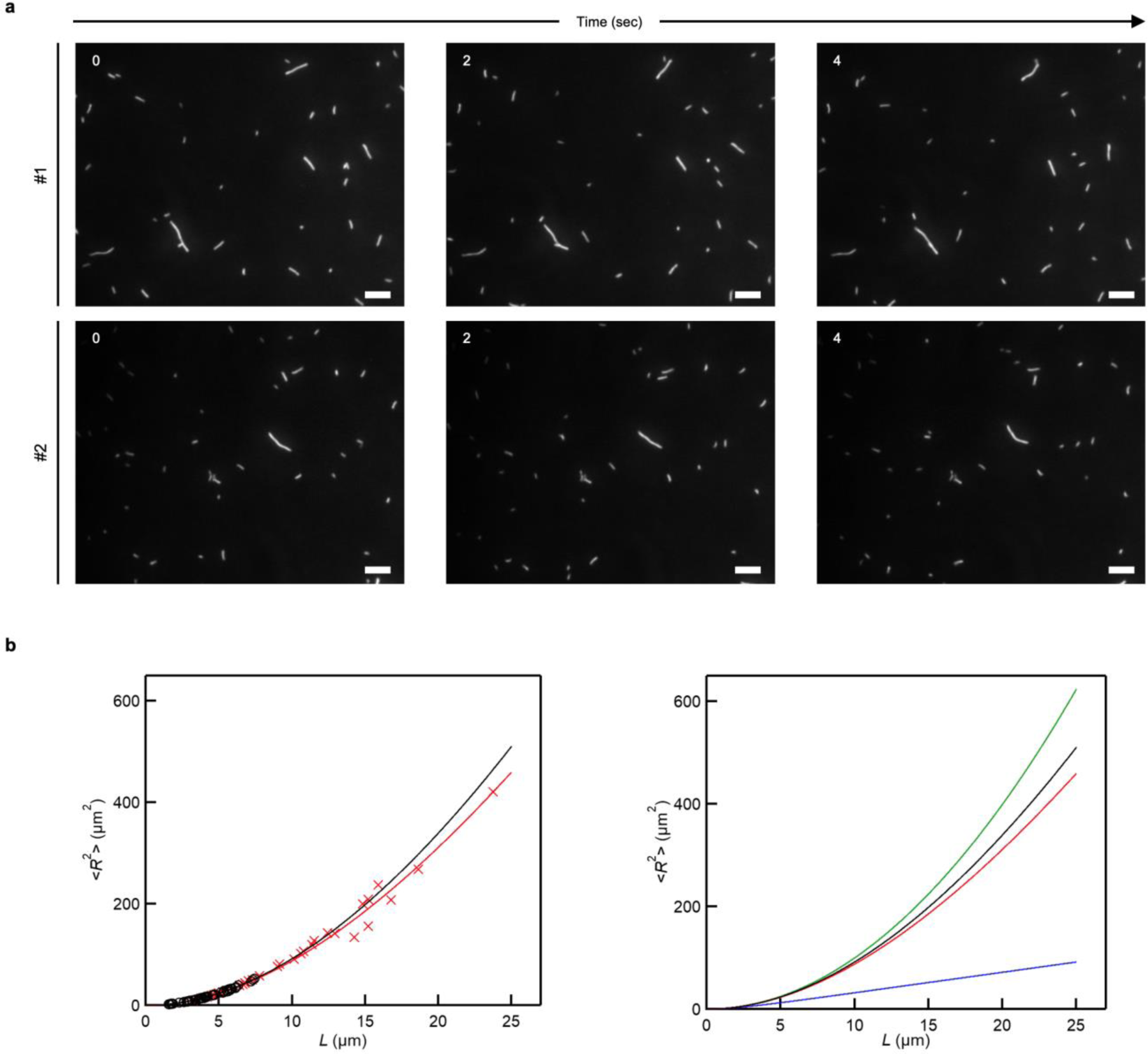
|Dynamic flexibility and persistence length analysis of PuuE tube. **a**, Time-lapse images of random bending of tube structures observed using TIRFM. Snapshots at the starting point (0 s) and after 2 and 4 s. Scale bars, 5 μm. **b,** Left, relationship between contour length (*L*) and mean square of end-to-end distance of the tube structures (<*R*^2^>). The continuous lines represent curves (black for PuuE tube, red for actin filament) fitted to the experimental data (black open circle for PuuE tube, red cross mark for actin filament). Wider range of *L* values than that in Fig. 3e were shown. Left, the theoretical curve of microtubules (green line) and that of intermediate filaments (blue line) are overlaid on PuuE tubes (black) and actin filaments (red). As *L* increases, the fitted curve of the tubes (black) increases similar to that of the actin filament but increases more slowly than the microtubule’s curve and faster than that of the intermediate filament.

**Extended Data Figure 9.**
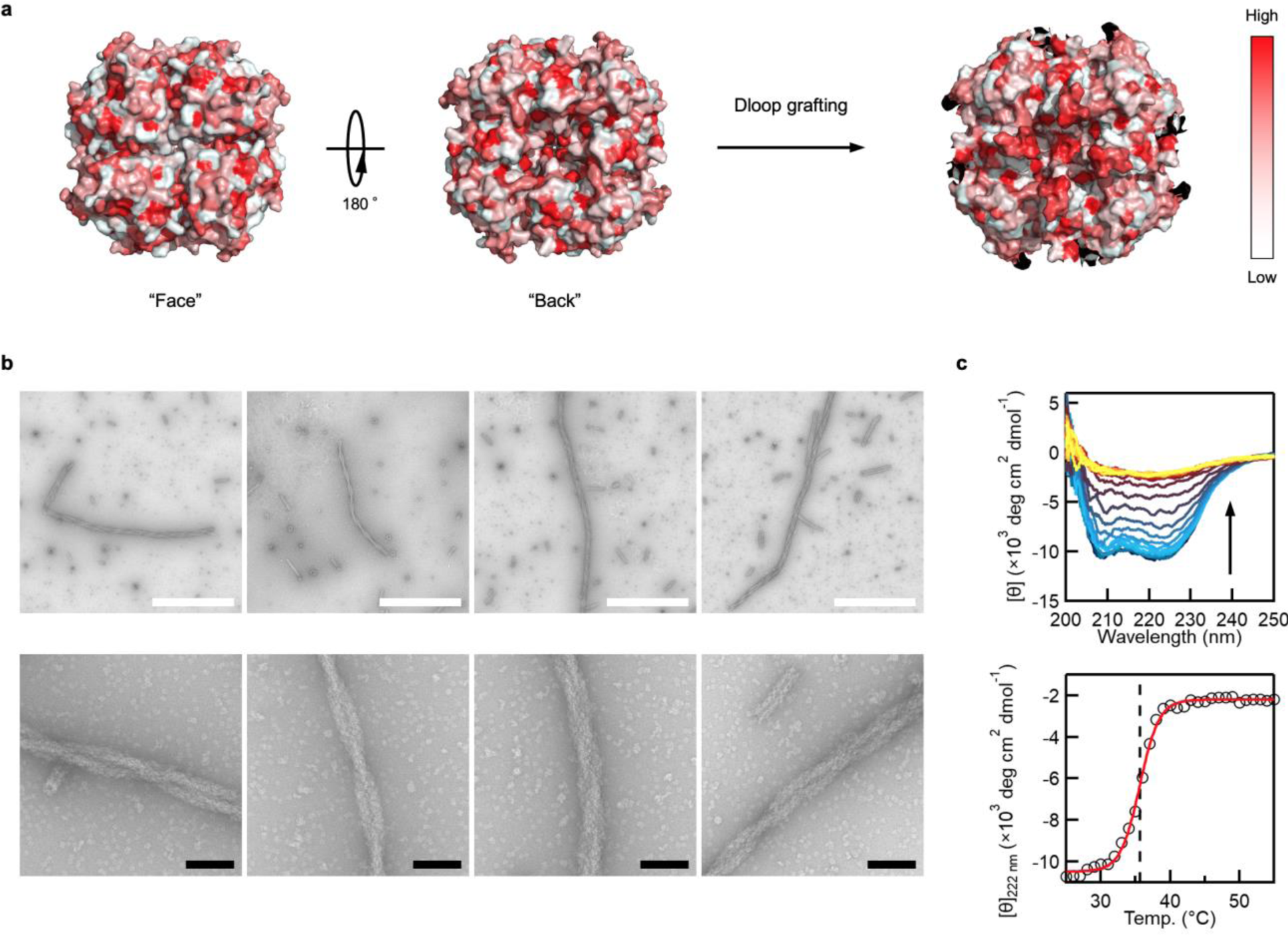
|D-loop grafting to emulate actin filaments. **a**, Surface hydrophobicity calculation for PuuE. D-loop was grafted into the “back” side of PuuE due to the hydrophobic nature of a prominent indentation. **b,** 12.5 μM of PuuE(D-loop)-M and PuuE-p in NaCl (+) buffer were incubated at 30 °C for 24 h and imaged using nsTEM. A novel helical pattern of two or three intertwined tubes was clearly observed. Flexibility was also noted when curved structures were observed. Scale bars, 1 μm (white), 100 nm (black). **c,** *T*_m_ measurement of PuuE(D-loop)-M via CD. 2.5 μM of PuuE(D-loop)-M in NaCl (+) buffer was incubated from 25 to 55 °C with temperature change of 1 °C/min. Top: overall CD spectra; bottom: thermal denaturation profiles, respectively.

**Extended Data Figure 10.**
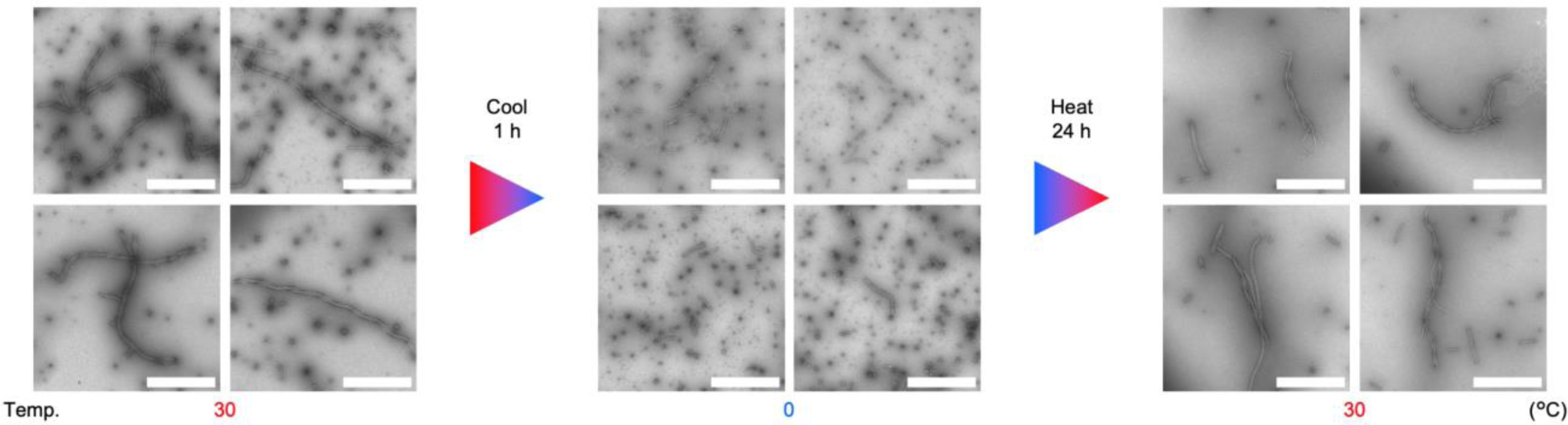
|Reversibility of structure formation in tubes with helical conformation. Additional images in Fig. 2d prove the reversibility of the tubular assemblies. For this analysis, we focused on the presence of helical tube structures. After 1 h at 0 °C, there were no such structures observed via nsTEM. Scale bars, 1 μm.

## Methods

### Plasmids and cloning

Primers for cloning and synthetic genes of N-terminal 6xHis-tagged PuuE-M and PuuE-p were purchased from Eurofins Genomics. PCRs were performed using the PrimeSTAR Max DNA Polymerase (Takara Bio) according to the manufacturer’s protocol. Sizes of PCR products were verified using standard agarose gel electrophoresis. The In-Fusion Snap Assembly (Takara Bio) was used as the standard method for cloning according to the manufacturer’s protocol, and each amplified gene fragment was ligated between the NdeI and BamHI multicloning sites of the pET11a expression vector (Novagen). Primers for cloning and a synthetic DNA fragment of Dloop were purchased from Eurofins Genomics. The plasmid encoding N-terminal 6xHis-tagged PuuE-Dloop-M was generated from the PuuE-M plasmid following the same procedures as above. All plasmids were amplified in *E. coli* strain DH5α (NIPPON GENE) and extracted using the NucleoSpin Plasmid EasyPure (MACHEREY-NAGEL) according to the manufacturer’s protocol. DNA sequences were confirmed by sequencing service (Eurofins Genomics).

### Protein expression and purification

The recombinant proteins were expressed using *E. coli* strain BL21 (DE3) (NIPPON GENE) co-transformed with a pGro7 chaperone plasmid (Takara Bio) and purified as follows. After transformation with plasmid DNA, colonies grown overnight on LB agar plates supplemented with 100 μg/mL ampicillin (Amp) and 20 μg/mL chloramphenicol (Crm) at 37 °C were picked to inoculate 5 mL of liquid LB-Amp-Crm broth and grown overnight at 37 °C and 200 rpm. Overnight cultures were diluted in 1 L of liquid LB-Amp-Crm broth supplemented with 0.5 mg/mL _L_-arabinose and grown at 37 °C and 200 rpm until reaching an optical density at 600 nm of 0.6-0.8. Protein synthesis was induced by adding 0.1 mM isopropyl-β-_D_-thiogalactopyranoside and the cultures were grown at 16 °C for 16-20 h. Cells were harvested by centrifugation at 10,000 rpm and 4 °C for 5 min and then frozen at -80 °C. Cell pellets were thawed at room temperature, resuspended in 60 mL of ice-cold purification buffer (20 mM Tris-HCl, pH 8.0, containing 300 mM NaCl) and lysed using sonication (9 min with 1:2 on/off cycles and 70% amplitude; SFX250, Branson) on ice. Cell debris was cleared by centrifugation at 10,000 rpm and 4 °C for 30 min. The supernatant (i.e., crude protein) was filtered through a 0.45 μm pore size membrane filter (Merck), applied onto HisTrap FF crude column (Cytiva) pre-equilibrated with the purification buffer and washed with 5 column volumes of 2% elution buffer (20 mM Tris-HCl, pH 8.0, containing 300 mM NaCl and 1 M imidazole; 2% means 20 mM imidazole). 6xHis-tagged proteins were eluted with 10 column volumes of elution buffer with a linear gradient of 2-40% (i.e., 20-400 mM imidazole). The fractions containing the proteins confirmed by means of UV absorption and SDS-PAGE were again collected and dialyzed against 50-fold volume of NaCl (+) or NaCl (-) buffer (50 mM Tris-HCl, pH 8.0, containing ±100 mM NaCl and 0.5 mM EDTA) at 4 °C twice. Each of the purified proteins was concentrated by an Amicon Ultra centrifugal filter units (Merck) with an appropriate molecular weight cutoff followed by filtration through a 0.45 μm pore size membrane filter (Merck). Protein concentration was determined by absorbance measurements at 280 nm using a NanoDrop OneC spectrophotometer (Thermo Scientific). The molar extinction coefficients at 280 nm for the proteins were calculated from the basis of amino acid composition^51^. The concentrated proteins were frozen in liquid nitrogen and stored at -80 °C prior to experiments.

### Sample preparation

All proteins were thawed immediately before tube formation experiments on ice. Each sample was prepared in a 1.5 mL microtube using appropriate buffer to adjust the concentration described in the manuscript and the volume to 200 μL at room temperature. Except for the NaCl concentration-dependent experiments, NaCl (+) protein stock solution and buffer were used. For the NaCl concentration-dependent experiments, 50 mM Tris-HCl (pH 8.0), 1 M NaCl and 0.5 mM EDTA was used in addition to NaCl (-) protein stock solution and buffer. Incubation of the samples was carried out using a ThermoMixer C (Eppendorf) or a MATRIX Orbital Delta Plus (IKA) with shaking of 300 rpm at the temperature described in the manuscript. For the disassembly and reassembly experiments, buffer substitution procedures were carried out using NaCl (-) and NaCl (+) buffer, respectively, with Microcon 50 centrifugal filter units (Merck) according to the manufacturer’s protocol four times at each step.

### Negative-stain transmission electron microscopy (nsTEM)

A naked G600TT copper grid (Nisshin EM) was carbon-coated using a VE-2030 (VACUUM DEVICE). The grid was glow-discharged using a PIB-10 (VACUUM DEVICE). Then, a 5-μL aliquot of the sample solution was placed on the grid for 1 min, and the remaining solution was removed with filter paper (No.2, ADVANTEC) followed by three-times rinsing with a 5-μL aliquot of Milli-Q water. After blotting off the water with filter paper, the sample was stained briefly with a 3-μL aliquot of 2% (w/v) uranyl acetate solution three-times. The remaining solution was removed with filter paper and the grid was dried on the bench-top. TEM observation was performed using a transmission electron microscope HT-7700 (Hitachi) with an acceleration voltage of 80 kV. The images were recorded using HT-7700 control software (Hitachi).

### Tube length analysis

Hundreds of discriminable tubes were picked-up manually on 5k-magnification TEM images. The tube lengths were calculated as half of the perimeter analyzed with ImageJ (Fiji)^52^. The plots were drawn by selecting 150 tubes from the longer lengths using Igor Pro 9 (WaveMetrics). For the disassembly and reassembly analysis, Welch’s t-test was carried out.

### Circular dichroism (CD) spectrum measurements

All proteins were thawed immediately before CD measurements on ice. Each sample was prepared in a 1.5 mL microtube using NaCl (+) buffer to adjust the concentration to 2.5 µM and the volume to 200 μL at room temperature. Far-UV CD spectra were obtained at a wavelength of 200-250 nm using a J-1100 spectropolarimeter (JASCO) with a quartz cell with light path of 1 mm. Thermal denaturation were performed at a temperature change rate of 1 °C/min. The CD spectral data were collected using Spectra Manager (version 2.5, JASCO). All CD data were expressed as mean residue ellipticity. The *T*_m_ of each protein was calculated from the thermal denaturation curve at wavelength of 222 nm by sigmoid fitting using Igor Pro 9 (WaveMetrics).

### Cryo-EM structural analysis

PuuE tube was diluted to one third of their original concentration. Quantifoil R1.2/1.3 Cu 300 grids coated with a holy carbon film (Quantifoil) were treated for hydrophilisation using a JEC-3000FC Auto Fine Coater (JEOL) at 20 Pa and 10 mA for 30 seconds. Subsequently, 2.5 µL aliquots of the respective diluted samples were applied to the prepared grids. After blotting off excess solution, the grids were rapidly immersed in liquid ethane for vitrification using a Vitrobot Mark IV (Thermo Fisher Scientific) at 4 °C and 100% humidity.

Sample screening and data acquisition were performed using a Glacios cryo-transmission electron microscope (Thermo Fisher Scientific) operated at an accelerated voltage of 200 kV, equipped with a Falcon4EC camera, at the Institute of Life and Medical Sciences, Kyoto University. Images were automatically acquired with the EPU software as movies with nominal magnifications and corresponding calibrated pixel sizes of 120,000x (1.22 Å/pixel).

### Cryo-EM image processing

Image analysis was conducted using the software package RELION 5.0beta^53,54^. A total of 4,346 movies were subjected to frame alignment using RELION’s own algorithm, followed by the contrast transfer function (CTF) was estimated with CTFFIND4^55^. Coordinates of the tubes were then manually registered, and 709,722 segments were extracted into 780×780-pixel boxes with an inter-box spacing of 30 Å. After five rounds of 2D classification, the segments were visually categorized by tube diameter in the 2D class-averaged images. These segments were then separately re-extracted and subjected to 3D refinement by applying *C*_4_, *C*_5_, and *C*_6_ symmetries, corresponding to increasing diameter. The classes with the fourth largest and larger diameters failed to produce reliable 3D structures due to significant heterogeneity. Subsequently, 3D classification without alignment was performed to analyse variations in tube diameters and helical patterns for the tubes with *C*_4_, *C*_5_, and *C*_6_ symmetries.

### Fluorescent labeling

Tube formation was carried out as described above under the optimised condition described in the manuscript. Labeling reaction was achieved by adding Alexa Fluor 488 succinimidyl ester dissolved in dimethyl sulfoxide (DMSO) to the tube solution at a final concentration of 0.7 mM. The reaction was then incubated at 25 °C for 1 hour with gentle shaking under shading. The excess dye was removed using NaCl (+) buffer with Microcon 300 centrifugal filter units (Merck) according to the manufacturer’s protocol four times. The labelled tubes were then stored under shading at 25 °C until further experiments.

### Fluorescence microscopy

An observation chamber was assembled by placing two double-sided tapes (thickness ∼100 µm) onto a silicone-coated coverslip (24 × 36 mm^2^, thickness No. 1; Matsunami) with another coverslip (18 × 18 mm^2^, thickness No. 1; Matsunami) on top. To passivate the surface of the coverslips against nonspecific adhesion of protein, the chamber was filled with 10 mg mL^−1^ of Pluronic F-127 (Sigma-Aldrich) dissolved in distilled water for more than 10 minutes at room temperature. After washing out Pluronic F-127 solution with 5 chamber volumes of NaCl (+) buffer, the chamber was filled with TIRFM buffer (50 mM Tris-HCl pH 8.0, 100 mM NaCl, 0.5 mM EDTA, 0.2%(w/v) methylcellulose (1500 cP, Wako), 1 mM DTT, 2 mM Trolox). Next, the Alexa488-tube solution was diluted to 1/10 in NaCl (+) solution, and further diluted to 1/10 (final 1/100 dilution) in TIRFM buffer. Then, the diluted tube solution was perfused into the observation chamber and sealed by Valap to prevent flow. The fluorescence images of tube structures were acquired at intervals of 40 ms with an inverted microscope (IX-71, Olympus) equipped with a 60× objective lens (PlanApo NA 1.45 oil, Olympus), an EMCCD camera (iXon3, Andor Technology) and an excitation laser with the wavelength at 488 nm (OBIS 488-60-LS, COHERENT). All observations were performed at 25 ± 1 °C.

### Mechanical property analysis

The persistence length of the tube structures was estimated as follows. First, the fluorescence images were converted to 8-bit images using ImageJ function. Then, the skeletons of the tube structures were tracked using ImageJ plugin, JFilament^56^. Distances between adjacent nodes composing the skeletons were set as 1 pixel. Next, the contour length (*L*) and end-to-end distance (*R*) of the tube structures at each frame were calculated using the coordinates of the nodes with custom-written python scripts. The mean square of *R* (〈*R*^2^〉) of each tube structure was calculated by averaging *R*^2^ along 100–200 frames.

〈*R*^2^〉 and *L* follow the following equation when the shape fluctuation is driven thermally^36^.

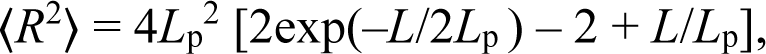

where *L*_p_ is the persistence length of the tube structure. The *L*_p_ values of the tube structures were estimated by fitting this equation to the experimental data using “curve_fit” function of Python package “scipy.optimize”. *L*_p_ of actin filaments was estimated by the same analysis. Totally, 55 tube structures and 37 actin filaments were analysed.

### Molecular Modelling

All predicted protein structures were generated by AlphaFold 2.2 or 2.3 multimer-mode (DeepMind)^15,16^. Cartoon models of the proteins were drawn using PyMOL 2.5 (Schrödinger)^57^ and UCSF ChimeraX (UCSF RBVI and NIH)^58^. Isoelectric points of PuuE-M and PuuE-p were calculated from the basis of amino acid composition^59^. Surface hydrophobicity of PuuE was drawn using Color_h script (PyMOL Wiki) based on the hydrophobicity scale^60^.

## Acknowledgements

This work was supported by JSPS KAKENHI (grant No. 19H02832, 19K22253 to Y.Suzuki, 21H05117 to Y.Suzuki and Y.Sugita, and 20K22628, 21J00530, 22KJ1644 to M.N.), JST PRESTO (grant No. JPMJPR22A7 to Y.Suzuki and JPMJPR20ED to M.M.), Takeda Science Foundation to Y.Suzuki, Chubei Itoh Foundation to Y.Suzuki, and The Hakubi Center for Advanced Research to Y.Sugita, M.M. and Y.Suzuki.

## Author contributions

Y.Suzuki directed the project. Y.Suzuki and M.N. conceived and designed the overall research. M.N. conducted experiment works with contributions from Y.Suzuki, Y.Sugita, and Y.Y.. Y.Sugita and M.N. performed cryo-EM data collection and analysed data. Y.Y. conducted TIRFM experiments, and Y.Y. and M.M. analysed mechanical property. M.N. and Y.Suzuki wrote a manuscript, with contributions from Y.Sugita, Y.Y. and M.M.

## Additional information

Supplementary Information. Amino acid sequence of proteins used in this study

## Supplementary Information

### Amino acid sequences

**PuuE-M;** 6xHis-TEVcs-PuuE-M3L2 MHHHHHHENLYFQGVDYPRDLIGYGSNPPHPHWPGKARIALSFVLNYEEGGERNILHG DKESEAFLSEMVSAQPLQGERNMSMESLYEYGSRAGVWRILKLFKAFDIPLTIFAVAMAA QRHPDVIRAMVAAGHEICSHGYRWIDYQYMDEAQEREHMLEAIRILTELTGERPLGWYT GRTGPNTRRLVMEEGGFLYDCDTYDDDLPYWEPNNPTGKPHLVIPYTLDTNDMRFTQV QGFNKGDDFFEYLKDAFDVLYAEGAEAPKMLSIGLHCRLIGRPARLAALQRFIEYAKSHE QVWFTRRVDIARHWHATHPYTSGQLVTPADIRRQARRVKKARERLAKALQADRLA

**PuuE-p;** 6xHis-TEVcs-PuuE-p66α MHHHHHHENLYFQGVDYPRDLIGYGSNPPHPHWPGKARIALSFVLNYEEGGERNILHG DKESEAFLSEMVSAQPLQGERNMSMESLYEYGSRAGVWRILKLFKAFDIPLTIFAVAMAA QRHPDVIRAMVAAGHEICSHGYRWIDYQYMDEAQEREHMLEAIRILTELTGERPLGWYT GRTGPNTRRLVMEEGGFLYDCDTYDDDLPYWEPNNPTGKPHLVIPYTLDTNDMRFTQV QGFNKGDDFFEYLKDAFDVLYAEGAEAPKMLSIGLHCRLIGRPARLAALQRFIEYAKSHE QVWFTRRVDIARHWHATHPYTPEERERMIKQLKEELRLEEAKLVLLKKLRQSQIQ

**PuuE(D-loop)-M;** 6xHis-TEVcs-PuuE(D-loop)-M3L2 MHHHHHHENLYFQGVDYPRDLIGYGSNPPHPHWPGKARIALSFVLNYEEGGERNILHG DKESEAFLSEMVSAQPLQQGVMVGMGQKGERNMSMESLYEYGSRAGVWRILKLFKAF DIPLTIFAVAMAAQRHPDVIRAMVAAGHEICSHGYRWIDYQYMDEAQEREHMLEAIRILT ELTGERPLGWYTGRTGPNTRRLVMEEGGFLYDCDTYDDDLPYWEPNNPTGKPHLVIPYT LDTNDMRFTQVQGFNKGDDFFEYLKDAFDVLYAEGAEAPKMLSIGLHCRLIGRPARLAA LQRFIEYAKSHEQVWFTRRVDIARHWHATHPYT SGQLVTPADIRRQARRVKKARERLAK ALQADRLA

## References

1 Zhu, J. et al. Protein Assembly by Design. Chem Rev 121, 13701–13796, doi:10.1021/acs.chemrev.1c00308 (2021).

2 Korn, E. D., Carlier, M. F. & Pantaloni, D. Actin polymerization and ATP hydrolysis. Science 238, 638–644, doi:10.1126/science.3672117 (1987).

3 Pollard, T. D. & Cooper, J. A. Actin, a central player in cell shape and movement. Science 326, 1208–1212, doi:10.1126/science.1175862 (2009).

4 Desai, A. & Mitchison, T. J. Microtubule polymerization dynamics. Annu Rev Cell Dev Biol 13, 83–117, doi:10.1146/annurev.cellbio.13.1.83 (1997).

5 Gudimchuk, N. B. & McIntosh, J. R. Regulation of microtubule dynamics, mechanics and function through the growing tip. Nat Rev Mol Cell Biol 22, 777–795, doi:10.1038/s41580-021-00399-x (2021).

6 Perlmutter, J. D. & Hagan, M. F. Mechanisms of virus assembly. Annu Rev Phys Chem 66, 217–239, doi:10.1146/annurev-physchem-040214-121637 (2015).

7 Sevvana, M., Klose, T. & Rossmann, M. G. Principles of Virus Structure. Encyclopedia of Virology, 257–277 (2021).

8 King, N. P. et al. Accurate design of co-assembling multi-component protein nanomaterials. Nature 510, 103–108, doi:10.1038/nature13404 (2014).

9 Bale, J. B. et al. Accurate design of megadalton-scale two-component icosahedral protein complexes. Science 353, 389–394, doi:10.1126/science.aaf8818 (2016).

10 Ben-Sasson, A. J. et al. Design of biologically active binary protein 2D materials. Nature 589, 468–473, doi:10.1038/s41586-020-03120-8 (2021).

11 Gnanapragasam, M. N. et al. p66Alpha-MBD2 coiled-coil interaction and recruitment of Mi-2 are critical for globin gene silencing by the MBD2-NuRD complex. Proc Natl Acad Sci U S A 108, 7487–7492, doi:10.1073/pnas.1015341108 (2011).

12 Walavalkar, N. M., Gordon, N. & Williams, D. C., Jr. Unique features of the anti-parallel, heterodimeric coiled-coil interaction between methyl-cytosine binding domain 2 (MBD2) homologues and GATA zinc finger domain containing 2A (GATAD2A/p66alpha). J Biol Chem 288, 3419–3427, doi:10.1074/jbc.M112.431346 (2013).

13 Suzuki, Y. et al. Self-assembly of coherently dynamic, auxetic, two-dimensional protein crystals. Nature 533, 369–373, doi:10.1038/nature17633 (2016).

14 Ramazzina, I. et al. Logical identification of an allantoinase analog (puuE) recruited from polysaccharide deacetylases. J Biol Chem 283, 23295–23304, doi:10.1074/jbc.M801195200 (2008).

15 Jumper, J. et al. Highly accurate protein structure prediction with AlphaFold. Nature 596, 583–589, doi:10.1038/s41586-021-03819-2 (2021).

16 Evans, R., et al. Protein complex prediction with AlphaFold-Multimer. bioRxiv, 2021.2010.2004.463034, doi:10.1101/2021.10.04.463034 (2022).

17 Patrian, M. et al. Supercharged Fluorescent Protein-Apoferritin Cocrystals for Lighting Applications. ACS Nano 17, 21206–21215, doi:10.1021/acsnano.3c05284 (2023).

18 Kang, H. et al. Identification of cation-binding sites on actin that drive polymerization and modulate bending stiffness. Proc Natl Acad Sci U S A 109, 16923–16927, doi:10.1073/pnas.1211078109 (2012).

19 Kang, H., Bradley, M. J., Elam, W. A. & De La Cruz, E. M. Regulation of actin by ion-linked equilibria. Biophys J 105, 2621–2628, doi:10.1016/j.bpj.2013.10.032 (2013).

20 Weisenberg, R. C. Microtubule formation in vitro in solutions containing low calcium concentrations. Science 177, 1104–1105, doi:10.1126/science.177.4054.1104 (1972).

21 Li, G. & Moore, J. K. Microtubule dynamics at low temperature: evidence that tubulin recycling limits assembly. Mol Biol Cell 31, 1154–1166, doi:10.1091/mbc.E19-11-0634 (2020).

22 Noji, M. et al. Heating during agitation of beta(2)-microglobulin reveals that supersaturation breakdown is required for amyloid fibril formation at neutral pH. J Biol Chem 294, 15826–15835, doi:10.1074/jbc.RA119.009971 (2019).

23 Goto, Y., Nakajima, K., Yamamoto, S. & Yamaguchi, K. Supersaturation, a Critical Factor Underlying Proteostasis of Amyloid Fibril Formation. J Mol Biol, 168475, doi:10.1016/j.jmb.2024.168475 (2024).

24 De Yoreo, J. J. et al. CRYSTAL GROWTH. Crystallization by particle attachment in synthetic, biogenic, and geologic environments. Science 349, aaa6760, doi:10.1126/science.aaa6760 (2015).

25 Chen, Y. et al. Morphology selection kinetics of crystallization in a sphere. Nature Physics 17, 121–127, doi:10.1038/s41567-020-0991-9 (2021).

26 Bruinsma, R. F., Wuite, G. J. L. & Roos, W. H. Physics of viral dynamics. Nat Rev Phys 3, 76–91, doi:10.1038/s42254-020-00267-1 (2021).

27 Kostiainen, M. A. et al. Electrostatic assembly of binary nanoparticle superlattices using protein cages. Nat Nanotechnol 8, 52–56, doi:10.1038/nnano.2012.220 (2013).

28 Liljestrom, V., Seitsonen, J. & Kostiainen, M. A. Electrostatic Self-Assembly of Soft Matter Nanoparticle Cocrystals with Tunable Lattice Parameters. ACS Nano 9, 11278–11285, doi:10.1021/acsnano.5b04912 (2015).

29 Anaya-Plaza, E. et al. Phthalocyanine-Virus Nanofibers as Heterogeneous Catalysts for Continuous-Flow Photo-Oxidation Processes. Adv Mater 31, e1902582, doi:10.1002/adma.201902582 (2019).

30 Liu, Q. et al. Optically Controlled Construction of Three-Dimensional Protein Arrays. Angew Chem Int Ed Engl 62, e202303880, doi:10.1002/anie.202303880 (2023).

31 Chalfie, M. & Thomson, J. N. Structural and functional diversity in the neuronal microtubules of Caenorhabditis elegans. J Cell Biol 93, 15–23, doi:10.1083/jcb.93.1.15 (1982).

32 Amos, L. A. Microtubule structure and its stabilisation. Org Biomol Chem 2, 2153–2160, doi:10.1039/b403634d (2004).

33 Chaaban, S. & Brouhard, G. J. A microtubule bestiary: structural diversity in tubulin polymers. Mol Biol Cell 28, 2924–2931, doi:10.1091/mbc.E16-05-0271 (2017).

34 Ferreira, J. L. et al. Variable microtubule architecture in the malaria parasite. Nat Commun 14, 1216, doi:10.1038/s41467-023-36627-5 (2023).

35 Golub, E. et al. Constructing protein polyhedra via orthogonal chemical interactions. Nature 578, 172–176, doi:10.1038/s41586-019-1928-2 (2020).

36 Landau, L. D., Lifshits, E. M. & Pitaevskiĭ, L. P. Statistical physics. 3rd ed., rev. and enl. / by E.M. Lifshitz and L.P. Pitaevskii, Repr. with corrections edn, (Pergamon Press, 1993).

37 Isambert, H. et al. Flexibility of actin filaments derived from thermal fluctuations. Effect of bound nucleotide, phalloidin, and muscle regulatory proteins. J Biol Chem 270, 11437–11444, doi:10.1074/jbc.270.19.11437 (1995).

38 Janson, M. E. & Dogterom, M. A bending mode analysis for growing microtubules: evidence for a velocity-dependent rigidity. Biophys J 87, 2723–2736, doi:10.1529/biophysj.103.038877 (2004).

39 Pampaloni, F. et al. Thermal fluctuations of grafted microtubules provide evidence of a length-dependent persistence length. Proc Natl Acad Sci U S A 103, 10248–10253, doi:10.1073/pnas.0603931103 (2006).

40 Van den Heuvel, M. G., de Graaff, M. P. & Dekker, C. Microtubule curvatures under perpendicular electric forces reveal a low persistence length. Proc Natl Acad Sci U S A 105, 7941–7946, doi:10.1073/pnas.0704169105 (2008).

41 Block, J., Schroeder, V., Pawelzyk, P., Willenbacher, N. & Koster, S. Physical properties of cytoplasmic intermediate filaments. Biochim Biophys Acta 1853, 3053–3064, doi:10.1016/j.bbamcr.2015.05.009 (2015).

42 Oda, T., Iwasa, M., Aihara, T., Maeda, Y. & Narita, A. The nature of the globular-to fibrous-actin transition. Nature 457, 441–445, doi:10.1038/nature07685 (2009).

43 Murakami, K. et al. Structural basis for actin assembly, activation of ATP hydrolysis, and delayed phosphate release. Cell 143, 275–287, doi:10.1016/j.cell.2010.09.034 (2010).

44 Durer, Z. A. et al. Structural states and dynamics of the D-loop in actin. Biophys J 103, 930–939, doi:10.1016/j.bpj.2012.07.030 (2012).

45 Das, S. et al. D-loop Dynamics and Near-Atomic-Resolution Cryo-EM Structure of Phalloidin-Bound F-Actin. Structure 28, 586–593 e583, doi:10.1016/j.str.2020.04.004 (2020).

46 Pratt, L. R., Chaudhari, M. I. & Rempe, S. B. Statistical Analyses of Hydrophobic Interactions: A Mini-Review. J Phys Chem B 120, 6455–6460, doi:10.1021/acs.jpcb.6b04082 (2016).

47 Koga, K. & Yamamoto, N. Hydrophobicity Varying with Temperature, Pressure, and Salt Concentration. J Phys Chem B 122, 3655–3665, doi:10.1021/acs.jpcb.7b12193 (2018).

48 Sun, Q., Fu, Y. F. & Wang, W. Q. Temperature effects on hydrophobic interactions: Implications for protein unfolding. Chem Phys 559, doi:ARTN 111550 10.1016/j.chemphys.2022.111550 (2022).

49 Echandia, E. L. & Piezzi, R. S. Microtubules in the nerve fibers of the toad Bufo arenarum Hensel. Effect of low temperature on the sciatic nerve. J Cell Biol 39, 491–497, doi:10.1083/jcb.39.2.491 (1968).

50 Nakajima, K. et al. Macromolecular crowding and supersaturation protect hemodialysis patients from the onset of dialysis-related amyloidosis. Nat Commun 13, 5689, doi:10.1038/s41467-022-33247-3 (2022).

51 Pace, C. N., Vajdos, F., Fee, L., Grimsley, G. & Gray, T. How to measure and predict the molar absorption coefficient of a protein. Protein Sci 4, 2411–2423, doi:10.1002/pro.5560041120 (1995).

52 Schindelin, J. et al. Fiji: an open-source platform for biological-image analysis. Nat Methods 9, 676–682, doi:10.1038/nmeth.2019 (2012).

53 Scheres, S. H. A Bayesian view on cryo-EM structure determination. J Mol Biol 415, 406–418, doi:10.1016/j.jmb.2011.11.010 (2012).

54 He, S. & Scheres, S. H. W. Helical reconstruction in RELION. J Struct Biol 198, 163–176, doi:10.1016/j.jsb.2017.02.003 (2017).

55 Rohou, A. & Grigorieff, N. CTFFIND4: Fast and accurate defocus estimation from electron micrographs. J Struct Biol 192, 216–221, doi:10.1016/j.jsb.2015.08.008 (2015).

56 Smith, M. B. et al. Segmentation and tracking of cytoskeletal filaments using open active contours. Cytoskeleton (Hoboken) 67, 693–705, doi:10.1002/cm.20481 (2010).

57 Schrodinger, LLC. The PyMOL Molecular Graphics System, Version 2.5.

58 Meng, E. C. et al. UCSF ChimeraX: Tools for structure building and analysis. Protein Sci 32, e4792, doi:10.1002/pro.4792 (2023).

59 Bjellqvist, B. et al. The focusing positions of polypeptides in immobilized pH gradients can be predicted from their amino acid sequences. Electrophoresis 14, 1023–1031, doi:10.1002/elps.11501401163 (1993).

60 Eisenberg, D., Schwarz, E., Komaromy, M. & Wall, R. Analysis of membrane and surface protein sequences with the hydrophobic moment plot. J Mol Biol 179, 125–142, doi:10.1016/0022-2836(84)90309-7 (1984).

